# ALG-2 and Peflin Stimulate or Inffibit Copii Targeting and Secretion in Response to Calcium Signaling

**DOI:** 10.1101/2020.02.22.944264

**Authors:** John Sargeant, Danette Seiler, Tucker Costain, Corina Madreiter-Sokolowski, David E. Gordon, Andrew A. Peden, Roland Malli, Wolfgang F. Graier, Jesse C. Ray

## Abstract

ER-to-Golgi transport is the first step in the constitutive secretory pathway which, unlike regulated secretion, is believed to proceed non-stop regardless of Ca^2+^ flux. Rowever, here we demonstrate that penta-EF hand (PEF) proteins ALG-2 and peflin constitute a hetero-bifunctional COPII regulator that responds to Ca^2+^ signaling by adjusting the ER export rate of COPII-sorted cargos up or down by ~50%. At steady-state Ca^2+^, ALG-2/peflin hetero-complexes bind to ER exit sites (ERES) through the ALG-2 subunit to confer a low, buffered secretion rate, while peflin-lacking ALG-2 complexes markedly stimulate secretion. During Ca^2+^ signaling, ALG-2 complexes lacking peflin can either increase or decrease the secretion rate depending on signaling intensity and duration-phenomena that could contribute to cellular growth and intercellular communication, following secretory increases, or protection from excitotoxicity and infection following decreases. In epithelial normal rat kidney (NRK) cells, the Ca^2+^-mobilizing agonist ATP causes ALG-2 to depress ER export, while in neuroendocrine PC12 cells, Ca^2+^ mobilization by ATP results in ALG-2-dependent enhancement of secretion. Within the NRK cell model, distinct Ca^2+^ signaling patterns can produce opposing ALG-2-dependent effects on secretion. Mechanistically, ALG-2-dependent depression of secretion involves decreased COPTT outer shell and increased peflin targeting to ERES, while ALG-2-dependent enhancement of secretion involves increased COPTT outer shell and decreased peflin at ERES. PEF protein complexes represent a true regulator of transport as they are dispensable for secretion yet adjust the secretion rate to physiological conditions. Their dynamics affects secretion of important physiological cargoes such as collagen T and significantly impacts ER stress.

## INTRODUCTION

The ER-to-Golgi interface is the busiest vesicle trafficking step, transporting up to one-third of all eukaryotic proteins (1). Anterograde cargo is captured into a COPTT pre-budding complex containing the inner coat sec23/24 heterodimer, which binds cargo in several distinct pockets on the membrane-proximal surface of sec24 (2–5). Recruitment of the outer coat layer, comprised of sec13/31, positions a flexible proline rich region (PRR) loop of sec31 across the membrane-distal surface of sec23, potentiating its Sar1 GAP activity required for cargo concentration (6). Together, the inner sec23/24 and outer sec13/31 COPTT coat involves polymerization of at least 24 hetero-tetramers (4).

Regulatory roles for Ca^2+^ in intracellular trafficking steps are still being elucidated. Recent work on ER-to-Golgi transport demonstrates a requirement for luminal Ca^2+^ stores at a stage following cargo folding/assembly, perhaps through the entry of Ca^2+^ into the cytoplasm where it binds and activates the vesicle budding, docking and/or fusion machinery (7, 8). Depletion of luminal calcium with sarco-endoplasmic reticulum Ca^2+^ ATPase (SERCA) inhibitors leads to significantly reduced transport as well as a buildup of budding and newly budded COPTT vesicles and vesicle proteins (7, 8). How Ca^2+^ causes these effects continues to be elusive, but part of the answer may lie with the penta-EF-hand-containing (PEF) protein adaptors that have been implicated in many Ca^2+^-dependent cellular phenomena (9). The PEF protein apoptosis-linked gene-2 (ALG-2) acts as a Ca^2+^ sensor at ER exit sites (ERES) and stabilizes association of sec31 with ERES through direct binding to a 12-amino acid sequence on the sec31A PRR region (10–13). Most ALG-2 in cell extracts exists in a stable heterodimer with the PEF protein peflin. Peflin binds ALG-2 in a Ca^2+^-inhibited manner (14, 15) and has been shown to suppress ER export of the cargo marker VSV-G-GFP, perhaps by modulating ALG-2 availability to bind ERES (16). Despite all of these observations, a unified model for when and how PEF proteins modulate secretion has not emerged. For example, most in vitro transport reconstitutions and results with purified ALG-2 have indicated that the protein is an inhibitor of vesicle budding or fusion (7, 17). On the other hand some recent intact cell trafficking experiments indicate a suppressive role for ALG-2 based upon ALG-2 depletion (18), while we implied a stimulatory role for ALG-2 because peflin suppressed transport by antagonizing stimulatory ALG-2-sec31A interactions (16). Furthermore, work on a presumed ALG-2 ortholog in yeast, Pef1p, demonstrated an inverse relationship, wherein Pef1p binding to the sec31 PRR was *inhibited* by Ca^2+^ and delayed coat recruitment to the membrane (19).

Here we advance understanding of PEF protein secretory regulation by demonstrating that ALG-2 binding to ERES can *either* stimulate *or* inhibit ER-to-Golgi transport depending upon ALG-2:peflin expression ratios and the nature of Ca^2+^ signals. Tn response to short bursts of agonist-driven Ca2+ signaling, ALG-2 increases outer coat targeting to ERES and stimulates transport. This response could help stimulated cells proliferate and/or replenish exhausted endocrine or exocrine secretory vesicles. On the other hand, a more relentless Ca^2+^ signal causes ALG-2 to markedly slow ER export. This novel physiological response that curtails COPTT targeting could represent a protective mechanism against excitotoxicity or infection.

## RESULTS

### Peflin expression levels determine ER-to-Golgi transport rates over a wide dynamic range in an ALG-2-dependent manner

To investigate the dynamic range and functional interactions of PEF protein regulation of ER export, we forced individual, tandem, or reciprocal expression changes of the two proteins. Endogenous peflin and ALG-2 were either knocked down using transfection with siRNA or over-expressed by transfection with the wt, untagged rodent proteins in NRK cells. After ≥24 hours of transfection, the initial rate of ER-to-Golgi transport of the synchronizeable transmembrane protein cargo VSV-G_ts045_-GFP was determined by incubation for 10 minutes at the permissive temperature followed immediately by fixation and morphological quantitation of the ratio of VSV-G that has reached the Golgi vs. remaining in the ER, as before (16). Figure 1 columns 1 and 2 show that as previously reported (16), peflin knockdown in the presence of normal levels of ALG-2 significantly increased VGV-G transport above basal by ~84%. On the other hand, over-expression of peflin (column 3) decreased transport by 23% below basal. Interestingly, the same two manipulations of ALG-2 expression in the presence of normal levels of peflin (columns 4 and 5) caused little change in transport relative to basal, indicating that at steady state, peflin expression levels are more rate-limiting. Forced peflin over- and under-expression thus defines a dynamic range of peflin regulation of transport at steady-state Ca^2+^ of ~107% of basal secretory flux (84% above basal and 23% below) in NRK cells. It is not known whether peflin expression ratios naturally vary substantially between different cell types or differentiation states.

**Figure 1.**
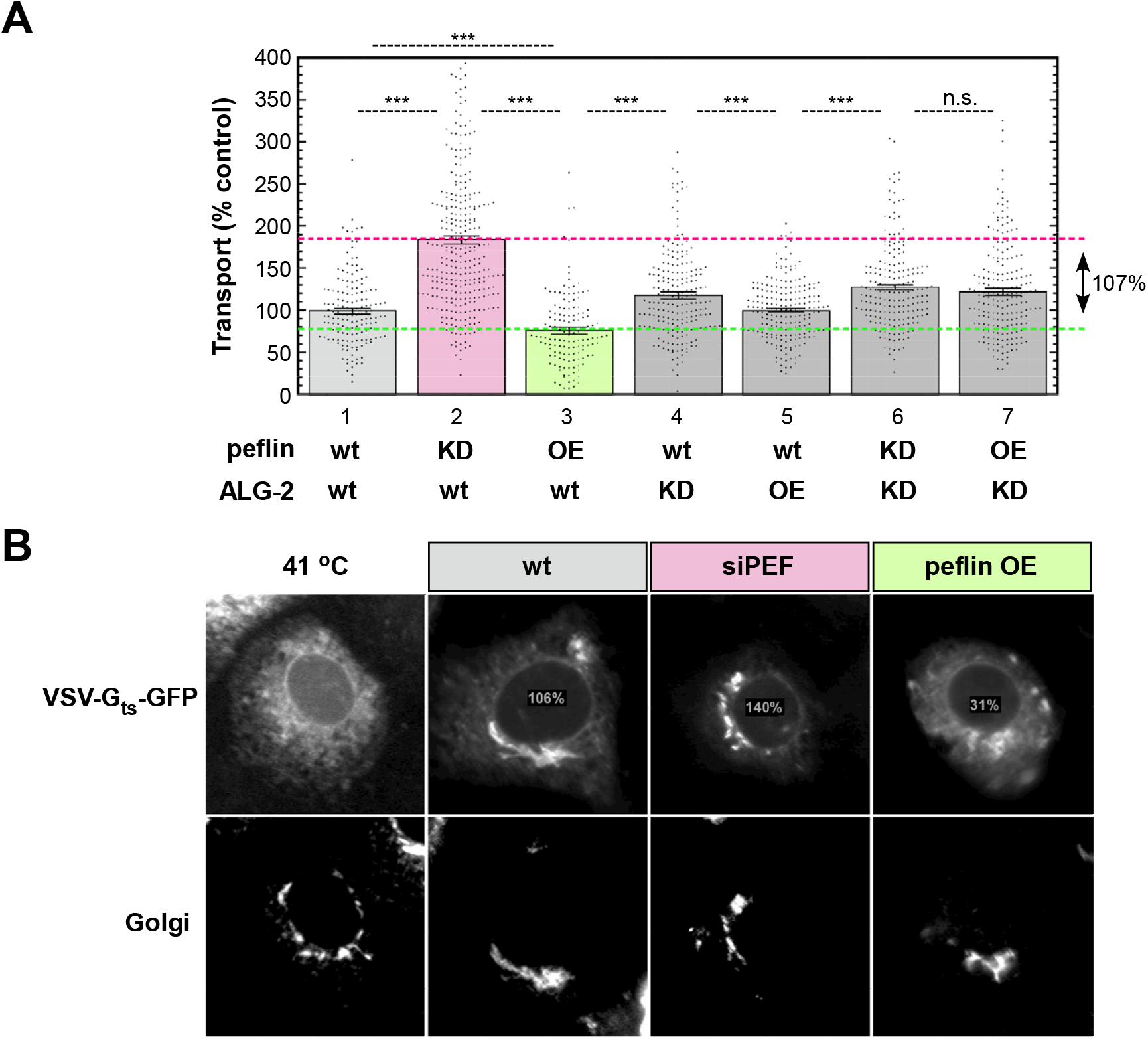
Peflin expression levels define a wide dynamic range of trafficking effects in an ALG-2 dependent manner. **(A)** NRK cells were transfected with VSV-G_ts045_-GFP with control or specific siRNAs and untagged over-expression constructs for peflin or ALG-2. Following growth at 41 °C, cells were shifted to 32 °C for 10 min to permit transport prior to fixation. Fixed cells were immuno-labeled with mannosidase II. Each transfected cell was assigned a transport index representing trafficking of VSV-G based upon the ratio of golgi intensity to peripheral ER fluorescence. The net transport index of each individual cell is plotted following subtraction of the mean transport index of cells kept at 41 °C, and normalization to the mean net transport index of wt cells. Approximately 200 cells were randomly quantified from each condition, and results shown are representative of at least 3 experiments with consistent trends. Asterisks indicate p values for unpaired T-tests for each set of conditions indicated with a dashed line. *, p<0.05; **, p<.005; ***, p<.0005. Standard error is shown for each plot. **(B)** Example widefield images of individual cells for select conditions with their transport index indicated as percent of control.

We next asked whether the effects of peflin over-expression and depletion depended upon the presence of ALG-2. As shown in Figure 1 columns 4 vs. 6 and 7, in ALG-2-depleted cells, forced changes in peflin expression do not change secretion, indicating that peflin is dependent upon ALG-2 to influence transport. This suggests that peflin’s effector for secretion is ALG-2. Interestingly, column 1 vs. 6 also indicates that in the absence of both ALG-2 and peflin, secretion is slightly higher than in the presence of both proteins at normal levels. This demonstrates that these PEF proteins are not required for transport and suggests that the two of them together exert a slightly suppressive effect on ER-to-Golgi transport under steady-state conditions.

### Peffin binds ERES via ALG-2 and prevents its stimufatory activity

Purification studies indicate that the majority of cellular ALG-2 is present in a 1:1 heterodimer with peflin (14). Since the subcellular distribution of peflin was unknown, we raised a rabbit polyclonal antibody against rat peflin to be used for localization studies of the endogenous protein. We also produced a chicken polyclonal antibody against mouse ALG-2 to be used in co-localization studies with peflin. Although peflin was previously thought to be a soluble cytoplasmic protein, we observed diffuse, reticular as well as distinctly punctate labeling for peflin throughout the cytoplasm (Figure 2C, upper right; also Figure 7A, upper row, third column). In addition, endogenous peflin was significantly concentrated in the nucleus. The labeling was specific for endogenous peflin since peflin siRNA transfection reduced all types of labeling (Figure 2C, third row, right column). Peflin cytosolic puncta noticeably co-localized with the ALG-2 cytosolic puncta previously identified as ER exit sites (ERES) (10–12), in these experiments also marked by sec13-GFP. Co-localization of endogenous peflin and ALG-2 at ERES was not expected, since previous research focused on peflin/ALG-2 heterodimers as a soluble species (14). We found that 95% of ERES defined by sec13-GFP were positive for ALG-2 and that the vast majority of ERES (>75%) were positive for both ALG-2 and peflin (supplemental Figure 1, columns 1 and 7). To determine the interdependence of peflin and ALG-2 for localization at ERES, we manipulated their expression levels as in Figure 1 and then quantified the labeling intensity of the two proteins specifically at ERES as defined by a sec13-GFP marker. Knockdown of ALG-2 removed peflin from ERES, implying that peflin was dependent on ALG-2 for targeting to ERES (not shown). Furthermore, in the absence of ALG-2, over-expression of peflin did not restore it to ERES (Figure 2A, bars 1 and 2), though peflin over-expression greatly increased peflin at ERES in the presence of ALG-2 (Figure 2A, column 4). ALG-2 targeting to ERES, on the other hand, did not depend upon, but yet was buffered by peflin. Peflin depletion greatly enhanced ALG-2 targeting to ERES (Figure 2B, columns 1 vs. 3), and peflin over-expression reduced it (Figure 2B bars 1 vs. 4). Targeting of the outer COPII coat subunit sec13-GFP (Figure 2D) mirrored the transport effects demonstrated in Figure 1 and extended the result we reported earlier for sec31A (16). That is, knockdown of peflin increased COPII targeting, potentially causing the observed increased transport, while peflin over-expression reduced COPII targeting, potentially inhibiting transport.

**Figure 2.**
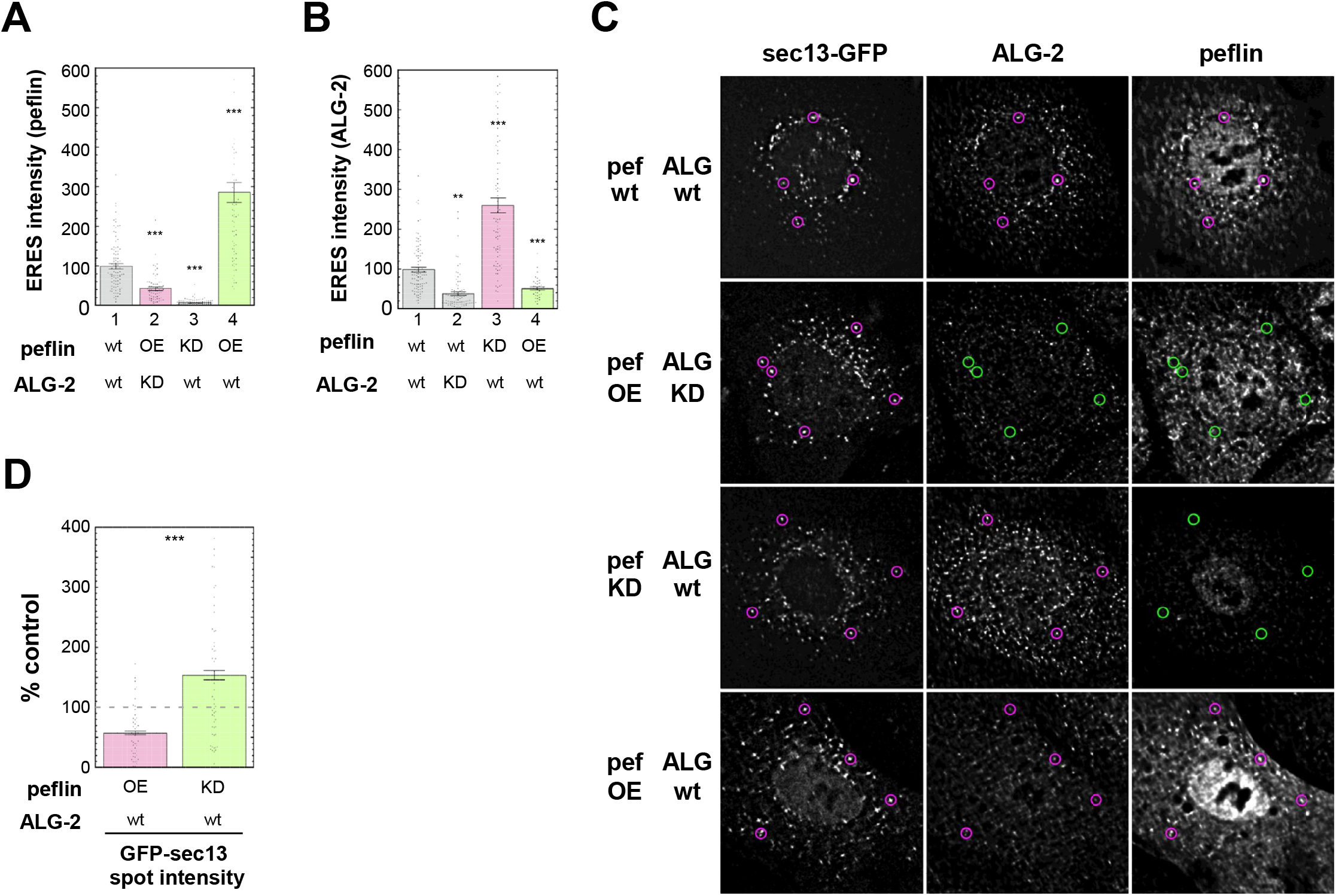
Peflin-ALG2 complexes localize to ER exit sites via ALG-2, competing with other ALG-2 complexes. NRK cells were transfected with GFP-sec13 and control or specific siRNAs and an untagged rat over-expression construct for peflin. Spots visible in the GFP-sec13 channel were defined as ERES. **(A)** Total intensity values of peflin colocalizing with ERES. Each point represents a single cell. Transfection conditions are specified below the graph and significance levels compared to column 1 are indicated above. Standard error is shown for each condition. **(B)** Total intensity values of ALG-2 colocalizing with ERES. **(C)** Deconvolved widefield immunofluorescence images of cells labelled with sec13-GFP, ALG-2 and peflin. Magenta circles highlight ERES containing peflin or ALG2 that co-localizes with GFP-sec13, while green circles note the absence of co-localization with GFP-sec13. Transfection conditions are specified to the left of images. **(D)** Total spot intensity of GFP sec13 in conditions with increased or decreased peflin expression levels, expressed as percent of control.

The targeting data indicates that peflin binds ERES through ALG-2 as part of an ALG-2-peflin complex, most likely the heterodimer species previously described (14). However, since removal of peflin increases ALG-2 at ERES, ALG-2 must also bind in other states, most likely the previously described homo-dimer for which a crystal structure is available with bound sec31A peptide (20). The high secretion caused by the lack of peflin and the low secretion caused by excess peflin are approximately equally above and below, respectively, the secretion in the absence of ALG-2 (Figure 1; height of column 4 is about mid-way between that of columns 2 and 3); this suggests that cytosolic peflin does not simply act as an ALG-2 sponge that decreases transport by withdrawing stimulatory ALG-2 from ERES - if this were the case, peflin over-expression could never inhibit transport below the level seen in the absence of ALG-2. Rather, peflin-containing ALG-2 species appear to themselves inhibit transport.

### ALG-2 can either stimulate or inhibit ER-to-Golgi transport, activities buffered by peflin

Figure 1A, columns 1 vs. 5, indicated that ALG-2 over-expression did not lead to significant changes in the ER export rate. This did not fit with other data. For example, since over-expression of ALG-2 should favor ALG-2 homomeric species over ALG-2-peflin hetero-complexes, as does a peflin knockdown, we wondered why ALG-2 over-expression did not stimulate transport. Thus, we titrated NRK cell transfections with 0.3, 1, and 3 micrograms of wildtype untagged ALG-2 construct DNA, and as shown in Figure 3A, black circles, these transfections resulted in 8-, 15- and ~25-fold over-expression of ALG-2 relative to control, demonstrating a titration of ALG-2 expression in individual cells. Figure 3B, black circles, then demonstrates that 8-fold over-expression stimulated transport by over 20%, 15-fold over-expression (the dose employed in Figure 1) had no significant effect, and ~25-fold over-expression inhibited transport by 20%. The same titration performed in the absence of peflin (magenta circles) resulted in similar biphasic effects, except that the inhibition was more potent and more severe relative to no over-expression (a drop from ~155% to ~95% of control). This experiment indicates that the relationship between ALG-2 and the secretion rate is complicated by several competing activities. First, it indicates that ALG-2 can both stimulate and inhibit transport, with stimulation giving way to inhibition as fold over-expression increases. Second, it indicates that there is both a stimulatory and inhibitory role for ALG-2 that are independent of peflin, in addition to the inhibitory role of ALG-2-peflin hetero-complexes implied in Figures 1–2. Third, it indicates that ALG-2-peflin hetero-complexes, though inhibitory, must buffer against the other inhibitory activity of ALG-2, revealed in this experiment, that is independent of peflin. Thus Figures 1–3, completed at steady-state Ca^2+^, suggest at least three distinct activities for ALG-2 and peflin complexes an inhibitory but buffering role for ALG-2-peflin hetero-complexes, a stimulatory role for ALG-2 without peflin, and an inhibitory role for ALG-2 without peflin.

**Figure 3.**
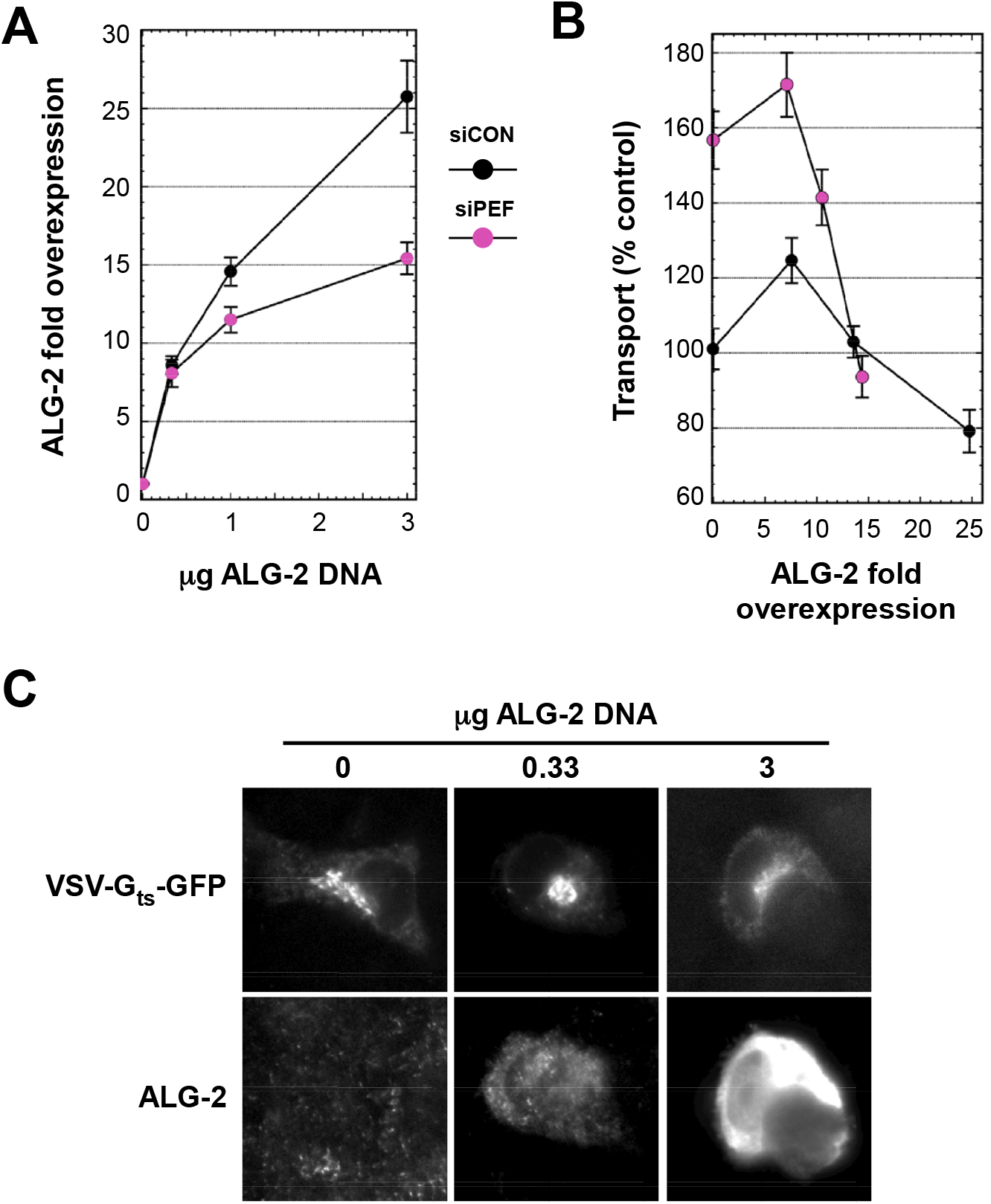
ALG-2 can inhibit or promote ER-to-Golgi transport independently of peflin. NRK cells transfected with control or peflin siRNAs were then re-transfected with VSV-G_ts045_ DNA and differing amounts of ALG-2 DNA. Each condition was assayed for mean ALG-2 florescence and accompanying transport index on a per cell basis. **(A)** Micrograms of ALG-2 DNA used in transfections versus fold overexpression determined by immunofluorescence quantitation. Fold overexpression for each cell was calculated as its own ALG-2 intensity divided by the mean intensity of untransfected cells. Mean ± SEM (n-100) is shown for each point. **(B)** The mean ER-to-Golgi transport value correlated to the mean ALG-2 fold overexpression level. (**C)** Example widefield images of individual siCON cells for select conditions.

### Peflin-ALG-2 complexes affect ER export similarly for multiple COPll client cargos, but not bulk flow cargo, and influence ER stress

A recent study reported that in an osteosarcoma cell line, peflin depletion inhibited ER-to-Golgi transport of collagen I, implying that peflin was required for collagen export from the ER (21). Since our results have instead suggested a suppressive role for peflin in VSV-G export, we investigated whether peflin may have opposite effects on different actively sorted cargoes. To address this, we expressed different cargoes in NRK cells and tested the effects of peflin depletion. These cargoes are schematized in Figure 4A. As seen if Figure 4B, compared to VSV-G-GFP, export of GFP-collagen I was even more strongly stimulated by peflin depletion (columns l and 2 vs. 3 and 4), supporting a suppressive effect of peflin under normal conditions. Since both VSV-G_ts045_-GFP and collagen export were synchronized by incubation at a restrictive temperature followed by a shift to permissive temperature, we wanted to rule out that the temperature shift was involved in the suppressive effects of peflin. We created novel reporter constructs containing a conditional aggregation domain, F_M_4, that aggregates in the ER and prevents export until a small molecule drug, AP2l998, is provided (22), causing synchronous ER export. The first construct included GFP as a luminal domain followed by F_M_4 and the VSV-G transmembrane domain (GFP-F_M_4-VSVG_tm_) which contains a di-acidic COPII sorting motif on the cytosolic surface. The second construct was similar but included the GPI anchor from CD55 at the C-terminus instead of a transmembrane domain (GFP-F_M_4-GPI). GPI anchors function as an export sequence recognized and sorted in a p24-dependent manner (23). Both constructs, when triggered by ligand AP2l998 were actively transported from the ER to the Golgi over a l0-minute time course. For both constructs, peflin depletion caused a highly significant increase in ER-to-Golgi transport (Figure 4, columns 5-8). A third construct, GFP-F_M_4-GH (24), was fully luminal, contained human growth hormone, and lacked any ER export sequence. Peflin depletions caused a significant *decrease* in ER export of this bulk flow construct. This result implies that peflin depletion does not act simply to accelerate vesicle production, but rather stimulates COPII function broadly--including its sorting function. It is also consistent with a recent study demonstrating that COPII sorting works in part by exclusion of proteins that are not actively included (25). In summary, Figure 4 establishes that the stimulatory effects of peflin depletion on transport are not restricted to high temperature-synchronized reporter cargoes, and that four actively sorted cargoes, VSV-G_ts045_-GFP, GFP-collagen I, GFP-F_M_4-VSV-G_tm_ and GFP-F_M_4-GPI, containing three distinct ER export signals - but not bulk flow cargo - are exported more efficiently in the absence of peflin in NRK cells. Thus peflin, it appears, is a bona fide suppressor of COPII function.

**Figure 4.**
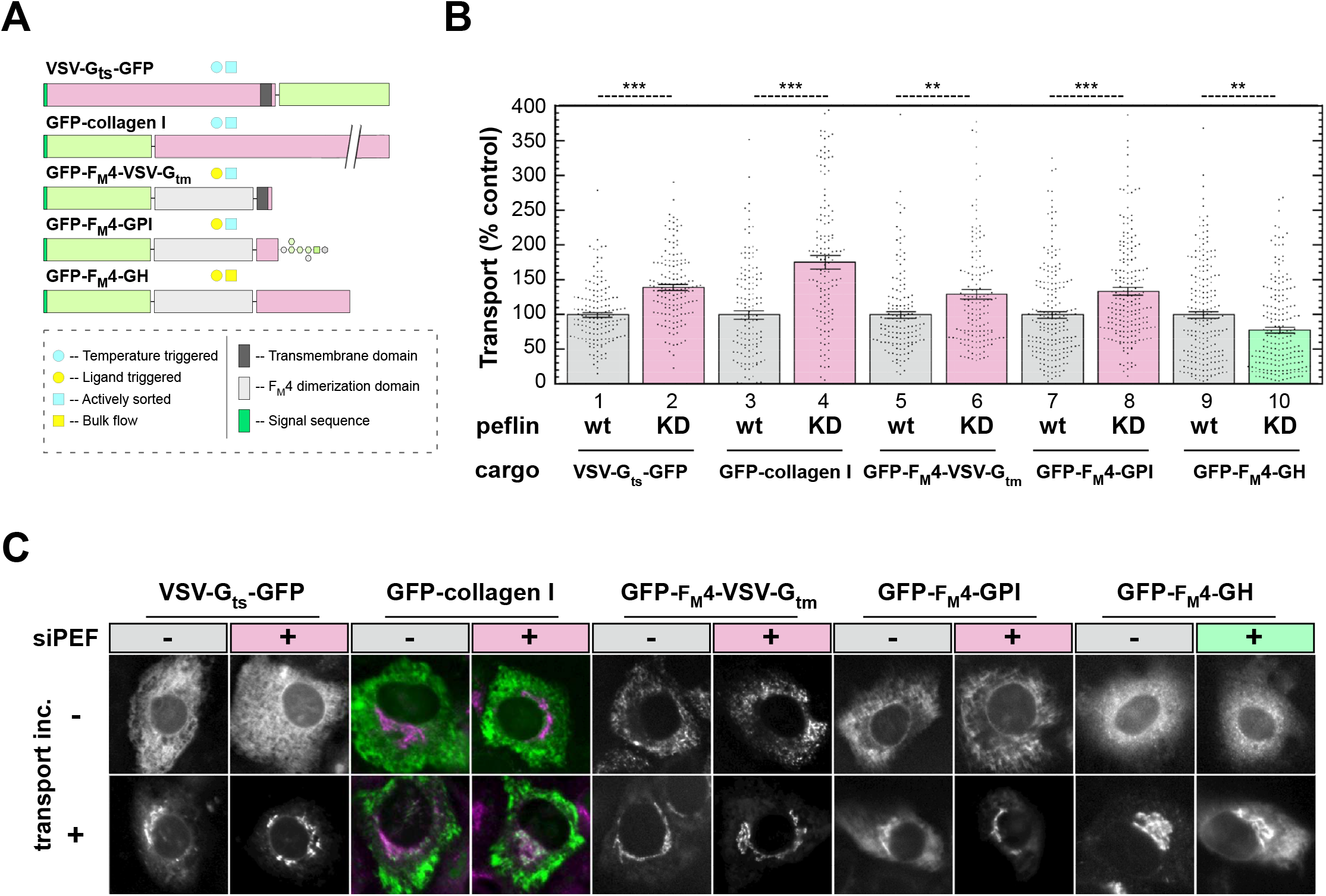
Peflin suppresses ER export of multiple actively exported cargos in NRK cells. **(A)** Schematic of cargo constructs used in B and C**. (B)** The initial rate of ER-to-Golgi transport was determined as in Figure 1 for NRK cells transfected with the indicated constructs in the presence of control or peflin siRNA. For VSV-G_ts045_-GFP and GFP-collagen I, transfected cells were placed at 41 °C to build up cargo in the ER. Transfer of cells to 32 °C provided a synchronous wave of transport to the Golgi. For GFP-F_M_4-VSV-G_tm_, GFP-F_M_4-GPI, and GFP-F_M_4-GH, cargo was accumulated in the ER under normal growth conditions at 37 °C, and transport was initiated by addition of the F_M_4-specific ligand. Mean ± SEM is shown for each condition, as well as approximate p values for the indicated unpaired T tests. **(C)** Example widefield images of individual cells for select conditions. For GFP-collagen I, a merge of GFP-collagen I and the Golgi marker Mannosidase II is shown since otherwise it was difficult to identify the Golgi following transport; all other images show the GFP channel only.

NRK cells may not be an adequate model for ER-to-Golgi transport for certain cargoes, for example collagen I, which require specific cargo adaptors and modified vesicles for efficient export (26–29). The vast majority of collagen I is secreted by fibroblasts, osteoblasts and chondrocytes. To address whether peflin also suppressed secretion of collagen I in cells whose normal function includes secretion of collagen I in abundance, we tested the effects of peflin depletion on endogenous collagen I secretion in Rat2 embryonic fibroblasts. The collagen I precursor, procollagen I folds inefficiently in the ER and mis-folded procollagen undergoes degradation by non-canonical autophagy at ERES (30). To be certain that non-secretory collagen fates potentially affected by peflin expression did not interfere with our assay for ER-to-Golgi transport, we monitored total cell fluorescence (TCF) of endogenous collagen I in addition to the ER-to-Golgi transport index. We measured ER-to-Golgi transport and collagen I TCF in the same cells with and without peflin depletion, and found that peflin depletion increased the ER-to-Golgi transport index by 75% (Figure 5A) but had no significant effect on collagen I TCF (Figure 5B). As shown in Figure 5C, collagen I TCF was quantitative and reflected collagen I content since titration of cells with a collagen I-specific siRNA resulted in distinct, decreasing TCF values. Together these data indicate that peflin depletion dramatically increases transport of endogenous collagen I from the ER to Golgi in fibroblasts and does not appear to affect collagen degradative pathways.

**Figure 5.**
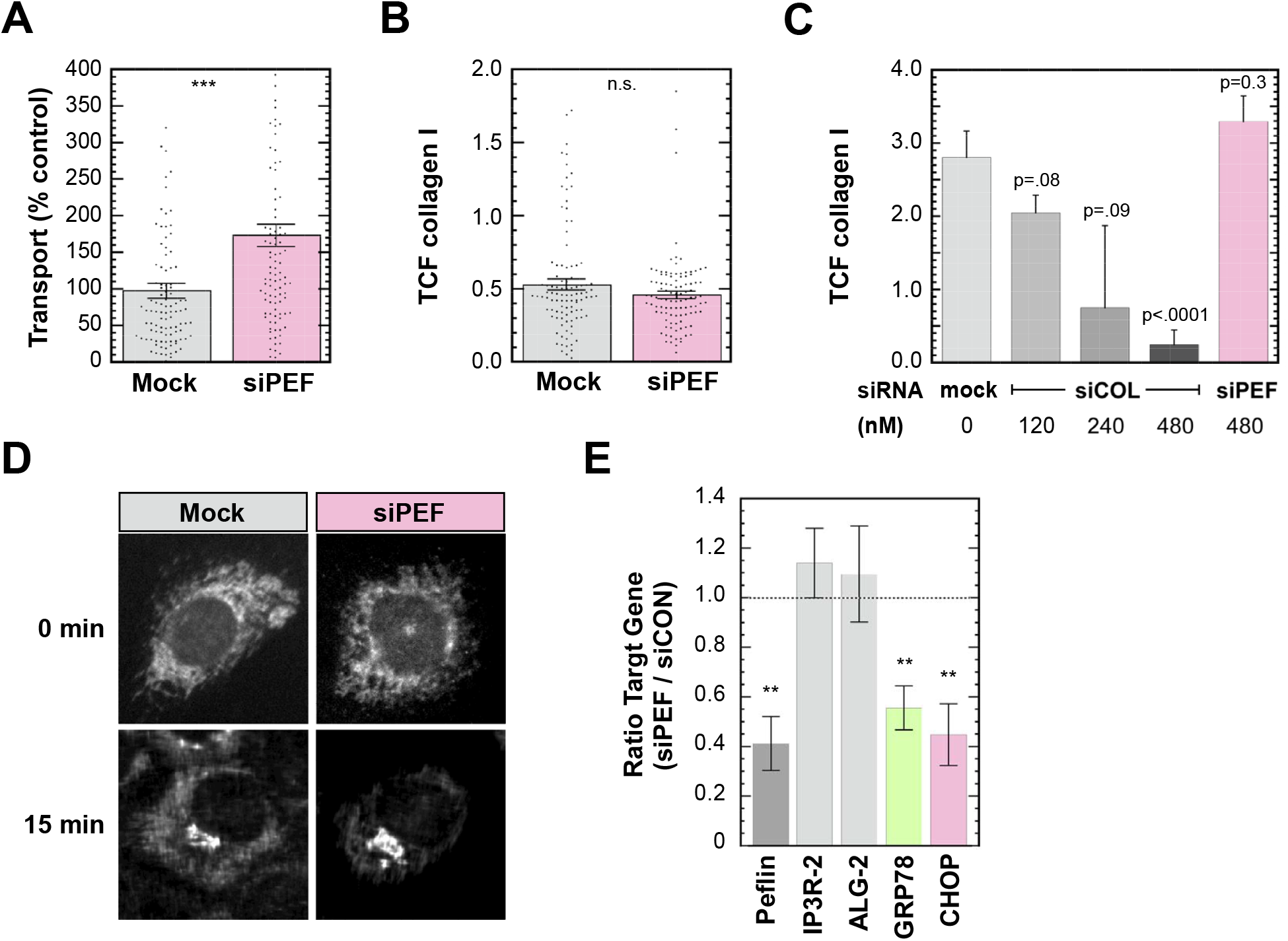
Peflin expression suppresses ER export of endogenous collagen I in Rat2 fibroblasts and facilitates pro-apoptotic UPR signaling in PAECs. Rat2 cells were transfected with the plasma membrane marker pCAG-mGFP in the presence or absence of peflin siRNA. Following growth at 41 °C for 24 h, cells were shifted to 32 °C with ascorbate-supplemented medium for 1S min to allow transport prior to fixation. **(A)** ER-to-Golgi transport assay employing collagen I immunofluorescence intensity in the ER and Golgi. **(B)** The same cells from A were analyzed for total cell fluorescence of collagen (see Experimental Procedures). **(C)** Validation of immunofluorescence assay for total cell fluorescence of collagen I. Cells were transfected with different concentrations of collagen I siRNA or an siRNA for peflin. Standard error and p values (vs. mock) is shown for each plot. **(D**) Example widefield images of collagen I immuno-labelling for select conditions in individual cells. **(E)** Peflin depletion in PS PAECs reduces GRP78 and DDIT expression. Senescent PS PAECs were subjected to control or siPeflin siRNA transfection and grown under standard conditions for 3 days prior to lysis and analysis by qRT-PCR for expression of several mRNAs as indicated beneath the plot. Results are shown as the ratio of mRNA expression in siPEF cells to that in siCon cells for each mRNA. Bars show mean ± SEM for 3 complete experiments conducted on different days using cells from different donors. p values indicate probabilities that the obtained ratios are equal to 1.0 (the null hypothesis).

So far we have shown that dramatically increasing the ALG-2 to peflin expression ratio can accelerate secretion of multiple cargoes. This suggests that decreasing peflin expression or another means of favoring the positive activity of ALG-2 could potentially relieve ER stress. To begin testing this idea, we utilized porcine aortic endothelial cells (PAECs), primary cells that undergo cellular ageing and senescence after passaging five times (P5) *in vitro*. As recently demonstrated (31), P5 PAECs display ER Ca^2+^-driven mitochondrial overload, oxidative stress, as well as profoundly increased unfolded protein response (UPR) signaling and expression of CHOP, a UPR transcription factor involved in the transition from UPR to apoptosis (32). Under these Ca^2+^ stress conditions, we found by quantitative reverse-transcription PCR (qRT-PCR) that a 60% knockdown of peflin, using siRNA, resulted in a specific 45% reduction in expression of the UPR target gene GRP78, and a 55% reduction of CHOP (Figure 5E). This demonstrates that peflin expression exacerbates life-threatening ER stress in aging endothelial cells, and that activation of secretion by ALG-2 can in principle relieve this burden.

### ln NRK cells, ALG-2 depresses ER export in response to sustained Ca^2+^ agonist stimulation

Since peflin and ALG-2 are regulated by Ca^2+^ binding in NRK cells, we tested whether their ability to regulate ER-to-Golgi transport was affected by cytoplasmic Ca^2+^ signaling. Histamine receptors present on many cell types activate phospholipase C via G_Q_ to stimulate Ca^2+^ release by IP3 receptor channels on the ER. As shown in Figure 6A (black circles), 10 minutes of ER-to-Golgi transport initiated after increasing times of exposure to histamine indicated that initially and for up to 30 minutes of exposure, no significant modulation of transport occurred. However, by 60 minutes of exposure, ER-to-Golgi transport was significantly reduced, with continued reduction for up to 150 minutes, wherein transport was reduced by 40% below basal. Thus, NRK epithelial cells respond to sustained Ca2+ agonist exposure by sharply curtailing ER secretory output, a new phenomenon we termed Ca^2+^-activated depression of ER export (CADEE).

**Figure 6.**
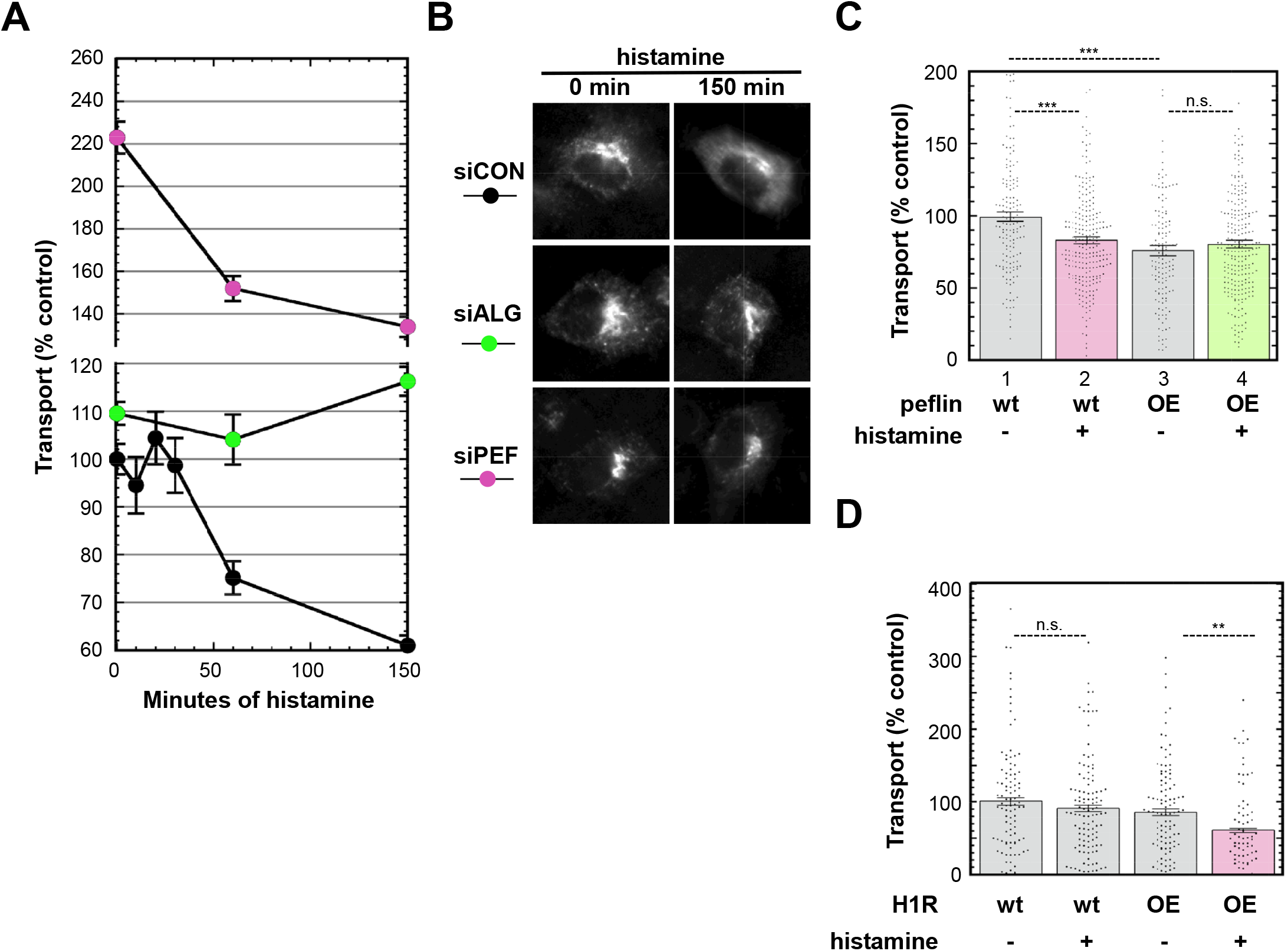
Ca^2+^-activated depression of ER export is mediated by ALG-2 independently of peflin. **(A)** NRK cells were transfected with VSV-G_ts045_-GFP along with control, ALG-2, or peflin siRNAs. Transfected cells were exposed to 100 μM histamine for 0-150 min at the non-permissive temperature prior to shift to the permissive temperature for 10 min, and transport was quantitated as in Figure 1. Mean ± SEM is shown for each point; n-150 cells per condition. **(B)** Example widefield images of individual cells for select conditions. **(C)** The rate of ER-to-Golgi transport following peflin overexpression and/or 100 μM histamine exposure for 150 min was determined as in Figure 1. **(D)** Transport index for NRK cells transfected with a control or H1R construct. These cells were exposed to +/- 50 μM histamine for 150 min.

We next tested the involvement of PEF proteins in the down-modulation. Significantly, the Ca^2+^-dependent modulation of transport was entirely dependent upon the presence of ALG-2, since knockdown of ALG-2 prevented any significant change in transport over the same timecourse (Figure 6A, green circles). The ALG-2-dependent activation mechanism, however, did not require peflin, since peflin knockdown did not prevent a histamine-activated decrease in ER-to-Golgi transport (Figure 6A, magenta circles). In the absence of peflin, however, ER export always remained well above control levels, indicating that although peflin is not the trigger, it still exerts a strong suppressive role throughout the Ca^2+^ signaling effect. Furthermore, Figure 6C demonstrates that over-expression of peflin inhibited transport (Figure 6C, columns 1 vs. 3), as shown earlier, but also protected against any inhibitory effects of histamine signaling (Figure 6C, columns 3 and 4). Together these results imply that the inhibitory effect of ALG-2-peflin hetero-complexes is an independent inhibitory state that can compete with a distinct, inhibitory ALG-2 activity during histamine signaling. One possibility is that histamine signaling activated the same inhibitory activity of ALG-2 observed when ALG-2 was over-expressed at steady-state Ca^2+^, which was also independent of peflin yet buffered against by ALG-2-peflin hetero-complexes (Figure 3).

While the histamine effect was repeatable numerous times, we found that NRK cells would occasionally respond not at all to histamine, a phenomenon that tended to occur several experiments in a row, perhaps indicating that specific lots of FBS or other unknown environmental factors might be affecting histamine responsiveness. This gave us the opportunity to directly assess the involvement of a single receptor system in the CADEE phenomenon. As shown in Figure 6D, in a single experiment NRK cells that were unable to modulate secretion in response to 50 uM histamine produced significant CADEE when transfected with the wildtype H1 histamine receptor, considered the main histamine receptor in kidney (33). This experiment defined a receptor system leading to CADEE in NRK cells.

### Sustained Ca^2+^ signaling decreases targeting of the COPll outer coat and increases targeting of peflin to ERES

To investigate the mechanism of the Ca^2+^-activated depression of ER export phenomenon, we monitored: outer coat subunits, peflin, ALG-2, and cargo at ERES by immunofluorescence microscopy in NRK cells with and without histamine treatment. Figure 7 parts A and B show representative images with different markers which, when quantitated revealed several significant changes. Most notably, at ERES containing ALG-2 and peflin, the outer coat labeling decreased in intensity. For example, using spots that contain ALG-2 and peflin to define ERES of interest, we found that GFP-sec13 intensity decreased by 40% at those ERES after histamine treatment (Figure 7C, left). This effect was due to a real change in sec13 intensity and was not a result of a change in the area of the regions of interest interrogated (Figure 7C, right). A similar 35% decrease was observed when measuring endogenous sec31A intensity using ALG-2 and the cargo VSV-G to define the measured ERES (Figure 7D), extending the trend to both subunits of the outer coat. If CADEE involved decreased targeting of outer coat to ERES by ALG-2, one prediction would be decreased co-localization of outer coat and ALG-2. Figure 7E demonstrates that there was a 40% decrease in GFP-sec13/ALG-2 overlap upon histamine treatment. Furthermore, the observed decrease in outer coat/ALG-2 overlap was not due to a detectable decrease in ALG-2 (data not shown), reinforcing the significance of decreased outer coat targeting (Figure 7C and D) as the driver of decreased co-localization. While the presence of ALG-2 under steady-state conditions has been implicated in stabilization of the outer coat (10, 13), our results may imply a role, under sustained Ca^2+^-signaling conditions, in which ALG-2 destabilizes the outer coat instead. Interestingly, despite the decrease in outer coat, we did not observe a decrease in GFP-sec13-peflin co-localization in the same cells (data not shown). This unexpected finding appeared to be due to a Ca^2+^-induced redistribution of peflin that counteracted the effects on co-localization of lost outer coat. Figure 7F, left, shows that cytoplasmic peflin spot intensity increased by 40% in the same cells in which the destabilized outer coat was documented. In these same cells, however, we detected a statistically significant 15% decrease in nuclear peflin spot intensity (Figure 7F, right). It is unknown whether peflin release from the nucleus is a passive result of, as opposed to a driver of, increased peflin targeting to ERES. Note that peflin, per se, is not required for CADEE induction but that during CADEE, peflin contributes substantially to the low transport values observed (Figure 6A). We currently do not know the significance of increased peflin at ERES during CADEE, but note that peflin-bound ALG-2 seems to be a parallel inhibitory state (Figure 6C). In conclusion, the CADEE phenomenon is accompanied by destabilization of the COPII outer coat at ERES and increased peflin localization at these sites, both predicted to decrease the rate of ER export.

**Figure 7.**
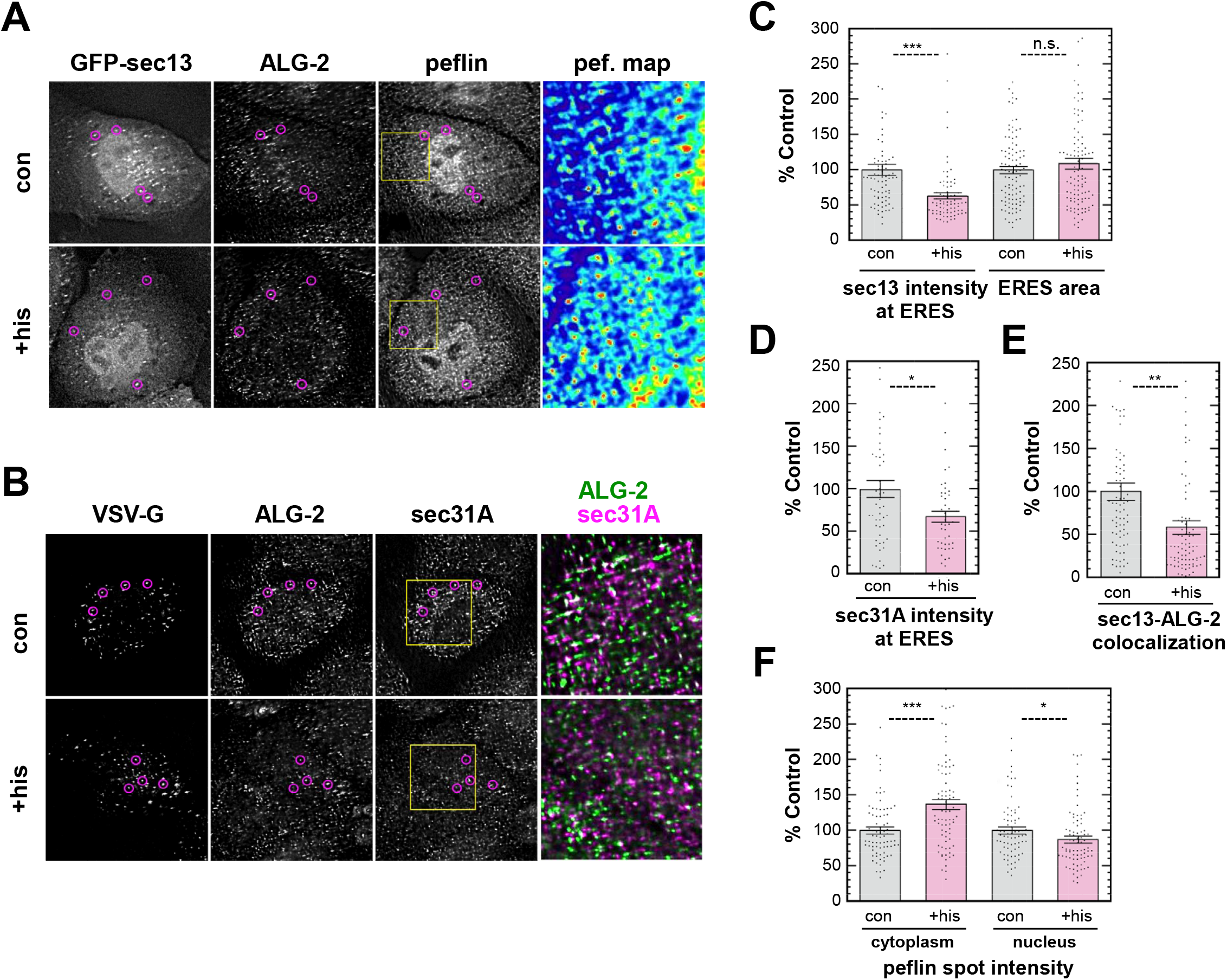
Histamine stimulation decreases outer COPTT coat and increases peflin targeting to ERES. **(A)** NRK cells were transfected with GFP-sec13, treated with or without histamine for 2.5 h, fixed and immuno-labeled for endogenous proteins. Shown are representative deconvolved images. Magenta circles mark several ERES positive for all three markers. GFP-sec13 intensity was lower at ERES in histamine-treated cells, while peflin intensity was higher. Peflin heatmap panels illustrate that there are more yellow/red objects in the cytoplasm following histamine treatment. **(B)** NRK cells were transfected with the transmembrane cargo VSG-G_ts045_, treated +/- histamine for 150 min at 41 °C, incubated for 45 s at 32 °C, fixed, immuno-labeled and displayed as in A. The conformation-specific VSV-G antibody used only recognizes assembled trimers. Sec31A intensity at ERES decreased upon histamine treatment, producing decreased co-localization with ALG-2, illustrated in the merged panels. **(C)** *Left*, an ROT was generated using ALG-2/peflin-colocalized spots to mark ERES and used to interrogate mean intensity in the unmodified GFP-sec13 channel. *Right*, the area of spots that had colocalized ALG-2 and peflin. **(D)** Using an ERES ROT generated from colocalized ALG-2 and VSV-G_ts045_ spots, total intensity of sec31A was measured. **(E)** Shows the total area of spots that had both GFP-sec13 and ALG-2. **(F)** *Left*, Peflin total spot intensity of cytoplasmic peflin. *Right*, the same calculation was performed for peflin particles inside the nucleus. Tn all plots, binary image masks were used to measure intensities on the unmodified images (see Experimental Procedures for details). Each dot represents data from a single cell, with the mean and SEM indicated.

### ATP elicits functionally distinct effects on ER export in NRK and PCJ2 cells

Extracellular ATP was used as a Ca^2+^ agonist and found to elicit CADEE similarly to histamine. A dose-response curve for ATP in ER-to-Golgi transport is shown in Figure SA. Interestingly, CADEE occurred maximally in doses close to 100 nM, while doses of 100 μM - a typical concentration in the literature for induction of Ca^2+^ signaling - resulted in little or no CADEE. This implied that the CADEE secretion response may depend upon on a defined Ca^2+^ signature or pattern during its induction. To investigate what the Ca^2+^ pattern associated with CADEE was, we recorded cytosolic Ca^2+^ signaling over a 40-minute period using FURA-2 cytosolic Ca^2+^ dye; cells were maintained at 37 °C in their regular medium during imaging. As shown in Figure SB, which displays representative individual cell traces, 100 nM ATP elicited a small, or sometimes no immediate Ca^2+^ response. During the subsequent 10 minutes following ATP addition, Ca^2+^ oscillations became more frequent and larger. Following a 15-minute break from imaging, cells with oscillations following ATP addition continued to show oscillations, now with even greater frequency and intensity. This experiment demonstrates that CADEE was associated with sustained Ca^2+^ oscillations over many minutes.

To examine the generality of these phenomena, we titrated neuroendocrine PC12 cells with the Ca^2+^ agonist ATP using the same cargo and ER-to-Golgi transport assay as in NRK cells. As shown in Figure 9A, PC12 cells responded to ATP with the opposite response to that of NRK cells, constituting a Ca^2+^-activated enhancement of ER export (CAEEE). The response to ATP involved an approximately 30% increase in ER export (as opposed to a 40% decrease in NRK cells) and was approximately 1 order of magnitude less sensitive to ATP, with a maximum near 1 uM ATP. Higher concentrations of ATP gave smaller effects on secretion as observed in NRK cells. To investigate the role of ALG-2 in the CAEEE response, we used siRNA to silence its expression. As shown in Figure 9B, PC12 cells depleted of ALG-2 did not change secretion in response to ATP, indicating that CAEEE, like CADEE, requires ALG-2. One possibility to explain the differing responses to ATP between NRK and PC12 cells would be that in different cell types, ALG-2 responds to Ca^2+^ in different ways because of differences in secretory machinery. Another possibility would be that ATP produces distinct Ca^2+^ signaling intensities or patterns in NRK and PC12 cells, and ALG-2 responds to distinct signaling properties in different ways. To help distinguish these possibilities, we carried out studies of cytoplasmic Ca^2+^ signaling in PC12 cells as we did in Figure 8 for NRK cells. As shown in Figure 9D, 1 μM ATP produced an immediate surge of cytoplasmic Ca^2+^ followed by a significant diminution within a few minutes. Thereafter, clear Ca^2+^ oscillations were rare and widely interspersed. These data suggest that distinct intensities and/or durations of Ca^2+^ signaling may underlie the different ALG-2-dependent secretory responses to ATP in NRK (Figure 8) and PC12 (Figure 9) cells. Sustained Ca^2+^ signals are associated with CADEE while strong responses followed by diminution are associated with CAEEE.

**Figure 8.**
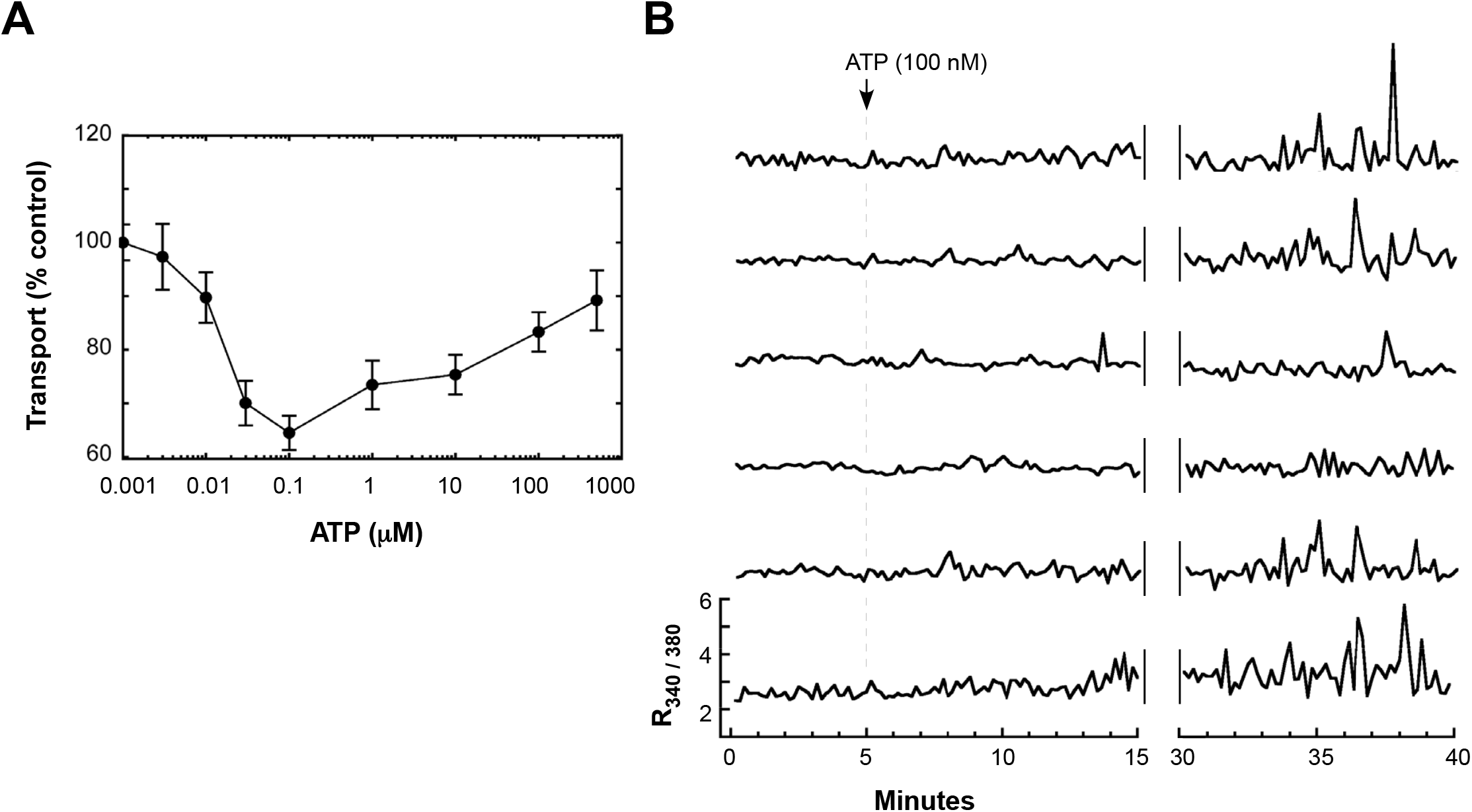
ATP elicits Ca^2+^-activated depression of ER export and continuing Ca^2+^ oscillations in NRK cells. **(A)** NRK cells were transfectd with VSV-G_ts045_ and treated with a range of ATP concentrations for 2.5 h at the non-permissive temperature prior to shift to the permissive temperature for 10 min, after which transport was quantitated as in Figure 1. Mean ± SEM is shown for each point; n-100 cells per condition. **(B)** Representative single cell FURA-2 traces. NRK cells were loaded with FURA-2 AM prior to incubation at 37 °C on the microscope stage in their regular medium supplemented with 20 mM Hepes. A first round of imaging every 10 s for 15 min was conducted, with introduction of ATP by pipet at t=5 min. At t=30 min, a second round of imaging every 10 s for 10 min was conducted on the same field of cells.

**Figure 9.**
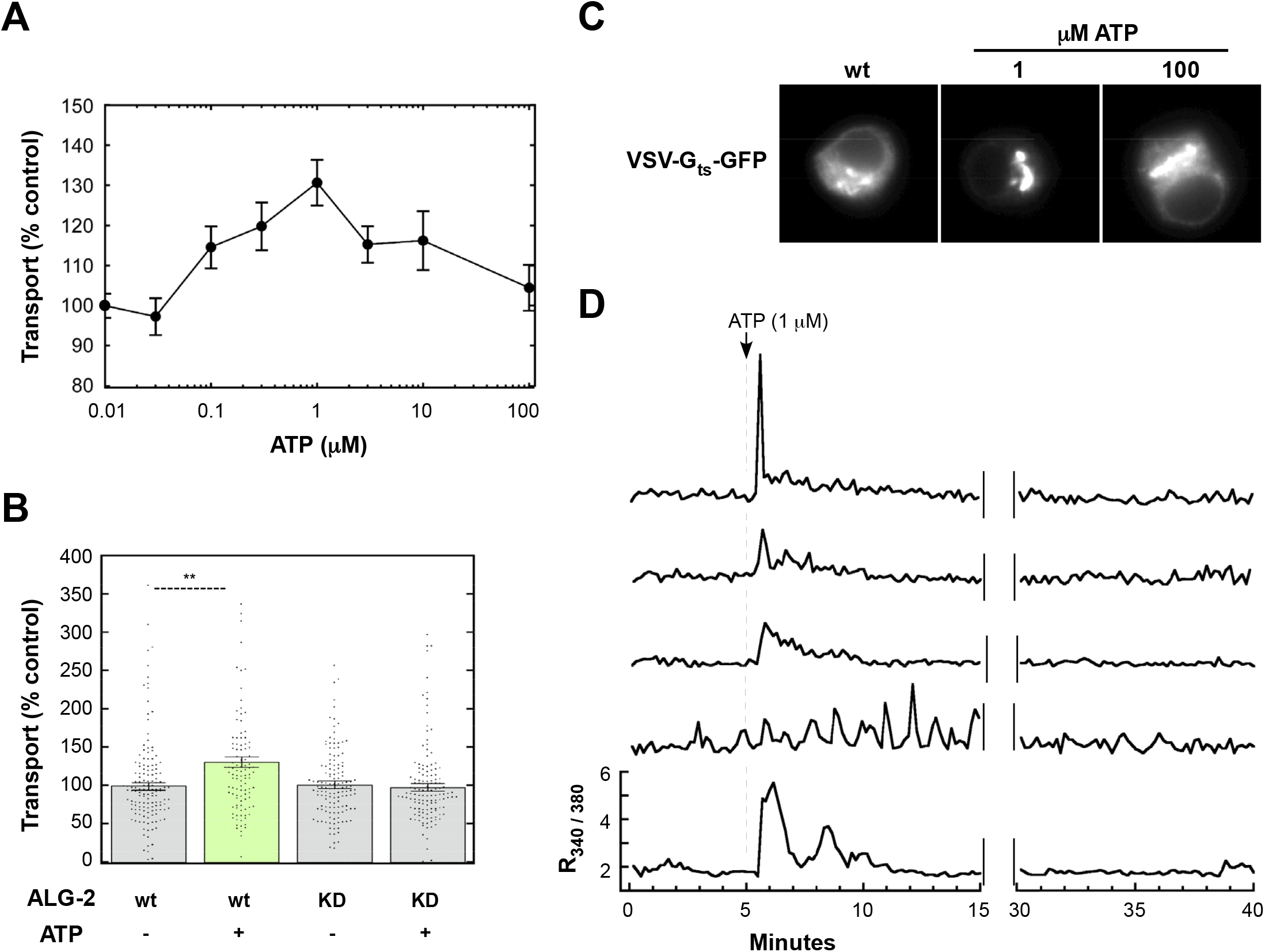
ATP elicits Ca^2+^-activated enhancement of ER export and discontinuous Ca^2+^ signaling in PCt2 cells. **(A)** PC12 cells were transfected with VSV-G_ts045_ and treated with a range of ATP concentrations for 2.5 h at the non-permissive temperature prior to shift to the permissive temperature for 10 min, after which transport was quantitated as in Figure 1. Mean ± SEM is shown for each point; n-100 cells per condition. **(B)** PC12 cells were transfected VSV-G and control or ALG-2 siRNA. Cells were exposed to 1 μM ATP for 2.5 h prior to the shift to a permissive trafficking temperature and quantitation of VSV-G transport. (**C)** Example widefield images of individual cells from A for select conditions. (**D)** In PC12 cells, exposure to 1 μM ATP produces immediate Ca^2+^ signaling that fades in strength and frequency over time. Shown are individual cell traces representative of this effect. The conditions used were identical to in Figure SB.

### ln NRK cells, distinct Ca^2+^ mobilization patterns determine whether ALG-2 enhances or depresses ER export

Since the opposing secretory responses of NRK and PC12 cells to ATP were associated with distinct Ca^2+^ signaling patterns, it seemed likely that spatiotemporal signaling properties, rather than a difference in secretory machinery, accounted for the distinct functional outcomes between the cell types. If this were the case, then NRK cells, given an appropriate Ca^2+^ stimulus, could respond with enhanced secretion like PC12 cells. Since an abrupt, discontinuous Ca^2+^ pattern was associated with CAEEE in PC12 cells (Figure 9), we tested the sarco-endoplasmic Ca^2+^ ATPase (SERCA) inhibitor 2,5-Di-(t-butyl)-1,4-hydroquinone (BHQ) in NRK cells, which invokes a surge of cytoplasmic Ca^2+^ for a few minutes until extrusion and other mechanisms restore the cytoplasm to near-basal levels (see below). We previously demonstrated that a combination of ATP and BHQ increased ER-to-Golgi transport of GFP-F_M_4-GH in HeLa cells, though a mechanism was not pursued further (34). As shown in Figure 10A, at low micromolar concentrations for 2.5 hours, BHQ stimulated ER-to-Golgi transport by 5O% or more, while by 10 μM, the compound caused no significant effect on transport (and in many cells, caused visible cytopathic effects). As shown in Figure 10B, the Ca^2+^-activated enhancement of ER export elicited by 4 μM BHQ was entirely ALG-2 dependent, since ALG-2 siRNA rendered cells unresponsive to BHQ. Representative individual cell traces of Ca^2+^ dynamics elicited by BHQ are presented in Figure 10C. BHQ produced a surge of Ca^2+^ that then diminished and stabilized. Compared to the 100 nM dose of ATP in NRK cells (Figure 8B) and the 1 μM dose of ATP in PC12 cells (Figure 9D), the 4 μM dose of BHQ more closely resembled the discontinuous pattern for ATP in PC12 cells that resulted in CAEEE. Together, these experiments indicate that ALG-2 responses to Ca^2+^ signaling can produce CADEE or CAEEE - even within the same cell type - based upon the signaling pattern and/or duration, with continuous, tonic elevations associated with the former and large surges followed by significant diminution associated with the latter.

**Figure 10.**
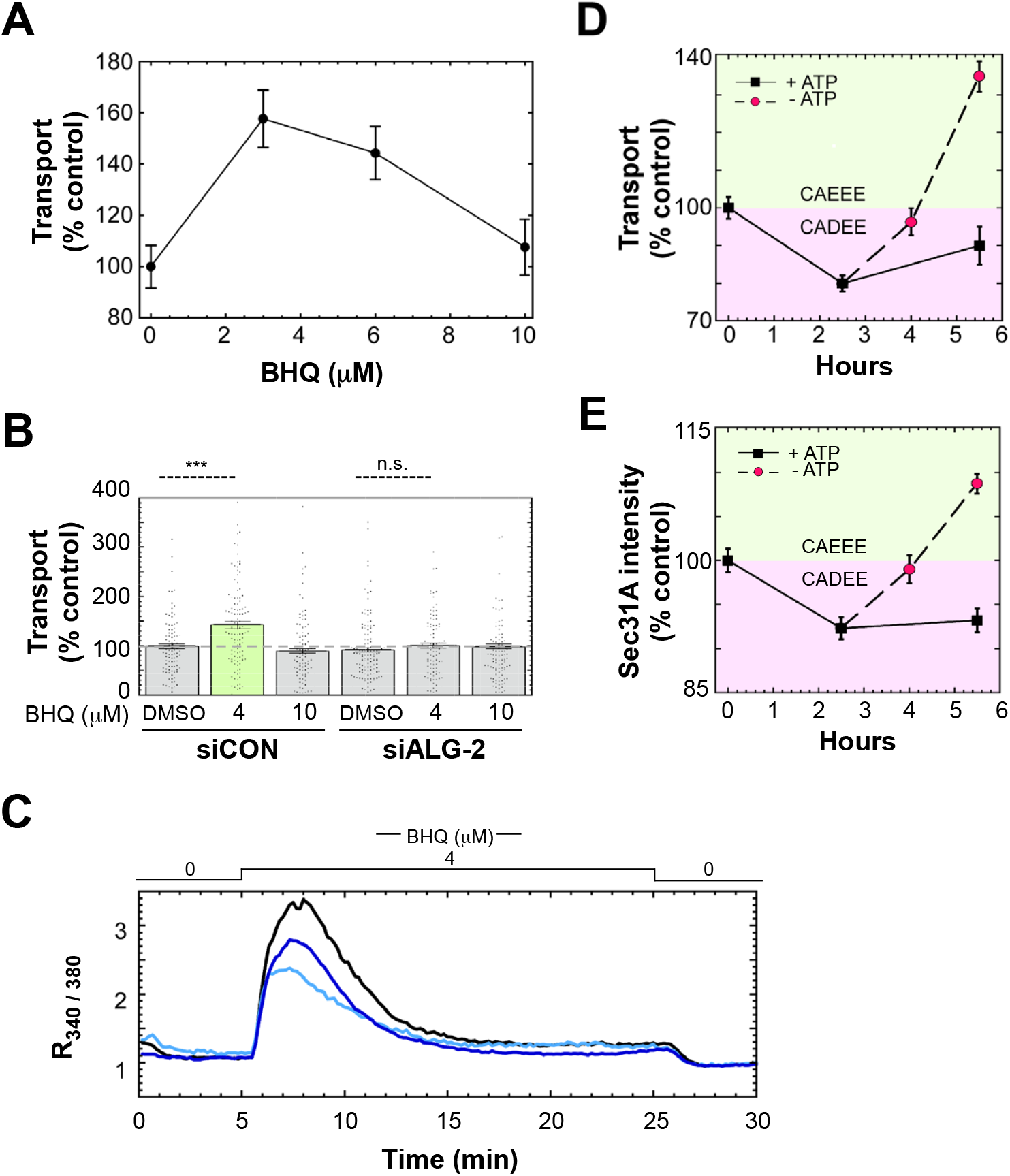
NRK cells possess both Ca^2+^-activated depression and enhancement of ER export, dependent upon the stimulation pattern and duration. **(A)** NRK cells transfected with VSV-G were exposed to a range of BHQ doses for 2.5 h after which transport indices were calculated for ~100 cells as for Figure 1. The mean transport index for each BHQ condition is plotted as a single black dot. **(B)** NRK cells transfected with VSV-G and control or ALG-2-specific siRNAs were treated with specific BHQ doses or a DMSO control for 2.5 h prior to determination of transport indices. **(C)** Example FURA-2 traces of NRK cells exposed to 4 μM BHQ. Methods similar to Figure 8B except that the cells were recorded during perfusion with media successively lacking, containing, and then lacking BHQ. **(D)** NRK cells were treated with 100 nM ATP and either maintained (black squares) for up to 5.5 h, or the ATP was removed by washing at 2.5 h and the cells were incubated for up to 3 h further (red circles). Transport relative to control was plotted for several timepoints. **(E)** Using cells from D, mean sec31 spot intensity was calculated from deconvolved images as in Figure 2D.

### ALG-2-dependent depression of transport is succeeded by the enhancement of transport upon agonist removal

Since both the ATP depressive effect and the BHQ enhancing effects in NRK cells depended upon Ca^2+^ and ALG-2, we doubted that they were mechanistically completely distinct. Instead, it seemed plausible that they were both part of a more uniform ALG-2 response that proceeded in consecutive phases, but that the relative intensity and timing of the phases could differ between different Ca^2+^ protocols or cell types. If this were the case, it should be possible to produce both CADEE and CAEEE using just ATP in NRK cells. Figure 10D black symbols shows ATP-dependent CADEE over 5.5 hours in NRK cells. As with the response to histamine (Figure 6), the CADEE effect of ATP is strong at 2.5 hours. After this initial induction, the depression may diminish somewhat but persists for an additional three hours. On the other hand as shown in the red symbols, if ATP is removed after the initial 2.5 hours, the secretion rate rebounds, achieving its initial rate 1.5 hours following ATP removal. Strikingly, by 3 hours following ATP removal (t = 5.5 h in Figure 10D), secretion has accelerated well beyond its initial, basal rate, an increase reminiscent of the enhancement of secretion caused by 2.5 hours of ATP treatment in PC12 cells (Figure 9A). While our results do not explain the molecular changes that result in the reversal of CADEE to cause CAEEE, it suggests that CADEE is not an off-pathway state but rather a state associated with ongoing signaling, while CAEEE may be a response to the cessation of signaling. In other words, both CADEE and CAEEE constitute a regulatory sequence controlled by the intensity and length of signaling episodes. This experiment using only ATP in NRK cells also rules out the possibility that potential signaling crosstalk between distinct signaling pathways determine whether CADEE or CAEEE manifests.

Given that changes in outer coat targeting to ERES would be a sufficient mechanistic basis for ALG-2- and peflin-dependent transport activities, we wondered whether the cycle of decreased and then increased transport induced by ATP incubation and removal (Figure 10D) would be reflected in parallel decreases followed by increases in outer coat targeting. Indeed, when sec31 spot intensity was quantified from the transport experiment shown in Figure 10D, we found that sec31A targeting closely correlated with CADEE, the release from CADEE by ATP removal, and subsequent induction of CAEEE by 5.5 hours (Figure 10E). These results support the hypothesis that ALG-2-dependent modulation of outer coat targeting drives all of the ALG-2-dependent transport phenotypes characterized herein.

## DISCUSSION

### ALG-2, peflin and effectors comprise a hetero-bifunctional regulator of ER export

From these studies, a model emerges (Figure 11) in which ALG-2, peflin and their effectors exist in several activity states correlating with distinct compositions of ER exit sites. These states result in either up-regulation (Figure 11B) or down-regulation (Figure 11C) of the basal ER export rate (Figure 11A) with potentially important consequences for cell physiology and pathology, including ER stress. The activity states are purely regulatory since neither peflin nor ALG-2 are essential for secretion. By experimentally adjusting the ALG-2 peflin ratio at steady-state Ca^2+^ we determined that a peflin-ALG-2 hetero-complex bound to ERES and inhibited ER-to-Golgi transport, and that a peflin-lacking ALG-2 species bound there to stimulate transport. Mechanistically, these states correlate with decreased and increased, respectively, targeting of the COPII outer coat to ERES, presumably mediated by the well-characterized interaction between ALG-2 and its binding site in the proline-rich region or loop of Sec31a that connects the inner and outer coats (2O). Though our studies did not define the subunit makeup or stoichiometry of ALG-2 complexes, it is likely that the inhibitory ALG-2-peflin complex targeted to ERES by ALG-2 is at least in part the 1 1 heterodimer characterized earlier to occupy the majority of ALG-2 in the cell and is dissociated by high Ca^2+^ (14, 15). The peflin-lacking ALG-2 complex is most likely comprised at least in part of the Ca^2+^-stimulated ALG-2 homodimer whose crystal structure was determined in complex with Sec31A peptide (20). Our schematic depicts them as a heterodimer and homodimer as the simplest explanation comprised of known species. The ALG-2 homodimer has been described generally as a Ca^2+^-dependent adaptor because of its ability to engage two co-effectors at once and link them (35). In our schematic, one of the ALG-2 co-effectors is always Sec31A, since that interaction is required for ALG-2 localization to ERES (10, 11). For the stimulatory activity of ALG-2 (Figure 11, part B), the simplest model would be that the other co-effector would be a second molecule of Sec31A, as this could result in cooperative coat assembly, more outer coat, and higher transport. Indeed, ALG-2-dependent inner-outer coat interactions have been demonstrated in solution without inclusion of other proteins (17). However, there may be other ALG-2 effectors involved that our work does not address, several of which have been identified at ERES, including annexin A11(18), MISSL, MAP1B (36), and KLHL12 (21).

**Figure 11.**
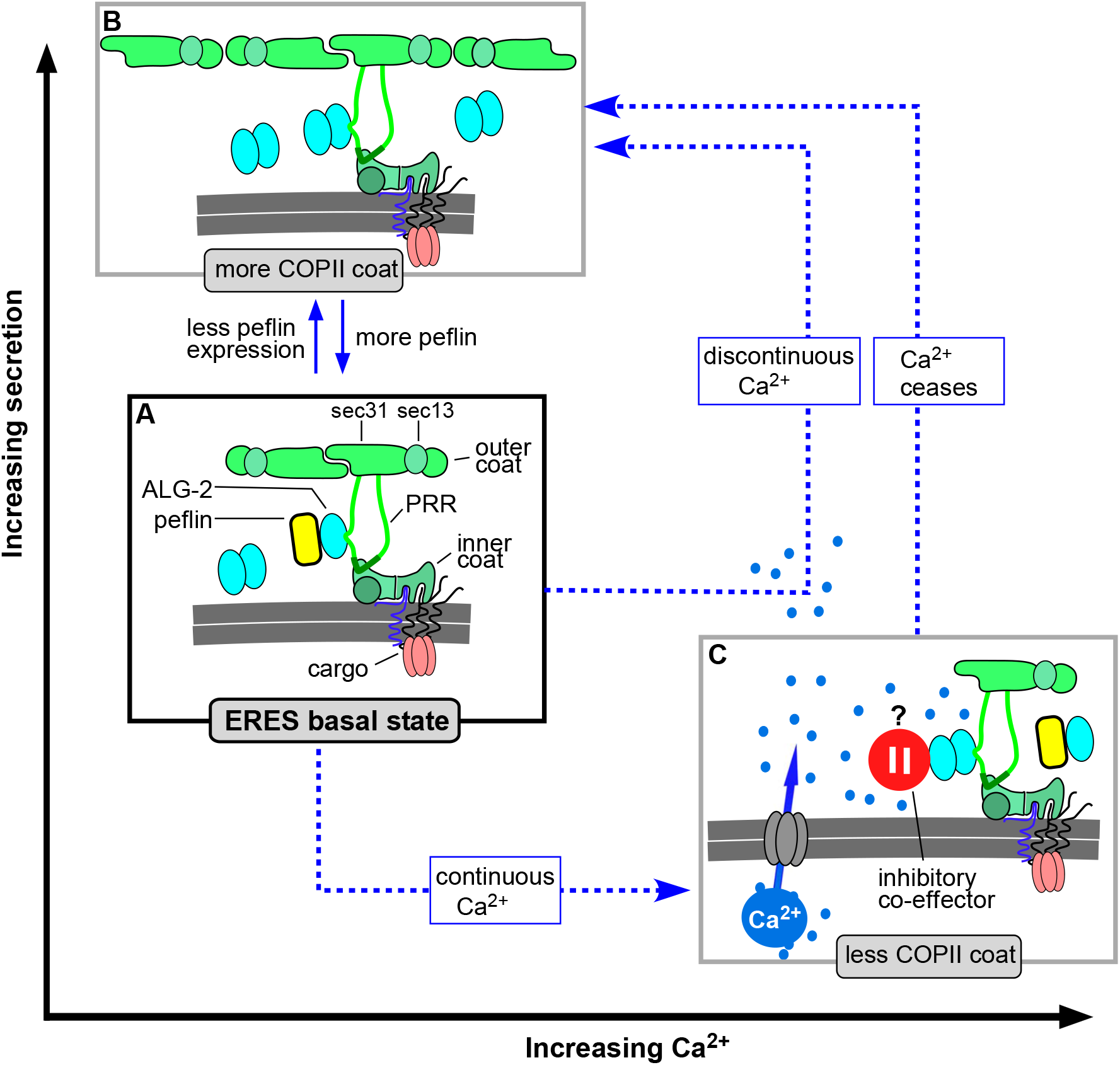
Model of PEF protein and Ca^2+^ regulation of ER export. **(A)** Under steady-state conditions, ALG-2 binds ERES in two distinct functional states that compete with each other. An ALG-2 homodimer binds sec31A to stimulate transport, while a peflin-ALG-2 complex binds sec31A via ALG-2 and inhibits transport. **(B)** Increases in the ALG-2:peflin expression ratio or discontinuous agonist-driven Ca^2+^ signaling comprised of high Ca^2+^ followed by low Ca^2+^, leads to more ALG-2 homodimers at ERES, greater outer coat recruitment, and greater export of COPII client cargo. **(C)** Continuous Ca^2+^ signals such as persistent agonist-driven oscillations leads to an ALG-2-dependent reduction of COPII targeting and decreased export of COPII client cargoes. Though peflin binding to ERES also increases during this state, it is not required for the response, leading us to posit either a distinct, unknown inhibitory Ca^2+^-activated ALG-2 effector (red pause button) or alternatively an inhibitory effect of ALG-2 produced by saturation of Sec31A (see Discussion). State C persists for hours if signaling persists, however, abrupt cessation of Ca^2+^ signaling leads directly to state B.

Ca^2+^ signaling can result in either a stimulatory or inhibitory activity of ALG-2, as can over-expression of ALG-2, that is independent of peflin. This stimulatory state is assumed in our model to be the same state just discussed and ascribed to the homodimer (Figure 11, part B). The Ca^2+^-induced inhibitory state could also be mediated by ALG-2 alone or could involve a distinct, inhibitory co-effector that is recruited by ALG-2 during sustained Ca^2+^ elevation (Figure 11, part C, red pause button). ALG-2 homodimers alone could be inhibitory because if the positive function of ALG-2 homodimers is to cross-link sec31A molecules, then over-saturation by homodimers should prevent that crosslinking. Though the cross-linking/over-saturation model to explain both influences is the simpler model, the distinct, inhibitory cofactor model actually fits our data better since it explains why depletion of ALG-2, or ALG-2 and peflin, does not per se inhibit secretion. We do not know what the putative inhibitory co-effector might be, but as just mentioned, ALG-2 has a number of Ca^2+^-dependent client proteins. By whichever mechanism, the inhibitory state is experimentally distinct from that containing peflin since it does not require peflin (Figures 3B, 6A) and is buffered against by the presence of peflin (Figures 3B, 6C).

Questions remain about how transitions occur between the identified activity states. For example, why does cessation of long-term Ca^2+^ stimulation lead to the stimulatory state (Figure 11, “Ca^2+^ ceases” blue arrow)? Perhaps during long-term Ca^2+^ signaling, ALG-2 homodimers accumulate at ERES, such that once the inhibitory co-effector is released, the stimulatory state mediated by homodimers automatically ensues. Another hole in our understanding is whether all Ca^2+^-induced secretion changes include at least a short, transient inhibitory state. An alternative possibility is that certain Ca^2+^ intensities and durations directly induce positive regulation of secretion (Figure 11 “discontinuous Ca^2+^” blue arrow). Unfortunately, this question is currently difficult to address given the lags before measurable secretion changes occur.

Another aspect to our model is that it only addresses the export of actively-sorted COPII client cargo. We have no data about whether vesicle budding rates, per se, increase and decrease in concert with the export rates of the measured cargos. It cannot be ruled out, for example that increases in VSV-G transport are actually caused by slower vesicle budding but more stringent sorting of client cargos. The fact that bulk flow cargo export slows as client cargo accelerates (Figure 4B) implies that client cargo sorting is so stringent as to exclude non-client cargo. What effect this has on vesicle budding remains unexplored.

An unexpected discovery was that peflin is concentrated in the nucleus (Figure 2). ALG-2 has also been reported to localize to the nucleus, and has been implicated in Ca^2+^-regulated splicing reactions there (37). While we cannot rule out the possibility that nuclear function of either ALG-2 or peflin could contribute to their transport effects, their presence and intensity ratios at ERES during expression studies (Figure 2) correlates extremely well with their functional impacts on ER-to-Golgi transport (Figure 1). We still do not understand, however, how Ca^2+^ changes result in the ERES targeting and functional changes we observed. For example, ALG-2 binding/unbinding to ERES occurs unitemporally with every single Ca^2+^ oscillation ((12), and our own unpublished observations), yet ALG-2-dependent changes in ERES structure/function caused by Ca^2+^ changes take ≥ 30 minutes. The unitemporal binding/unbinding may contribute to a longer-term change, perhaps involving the entire cellular balance of ALG-2 homodimers vs. heterodimers, that actually drives the functional changes. Alternatively, the ERES functional changes may involve something more akin to ERES biogenesis and disassembly, rather than merely changes in the content of outer coated located at ERES. One possibility we have largely ruled out involves the transcriptional response to ER stress. Knockdowns of peflin in unstressed cells under basal conditions did not detectably affect the unfolded protein response (UPR) as indicated by intensities of bands on Westerns with the following antibodies: anti-phospho-Ire1, anti-phosho-EIF2 alpha, anti-ATF4 and anti-CHOP (data not shown). This excludes the mechanism wherein peflin depletion could cause ER stress which would increase transcription of COPII machinery to build ERES and accelerate secretion. Based upon all available evidence, we conclude that PEF protein effects on ER-to-Golgi transport are mediated directly through, or triggered by, their interactions with sec31A at ERES.

There are many reasons for cells to up- or down-regulate ER export, including to regulate inter-cellular communication in the case of secretory cells and neurons, to regulate resource consumption, cell growth, ER stress, and to limit viral replication. It is of note that there appear to be two inhibitory states, one peflin-dependent and another peflin-independent. The peflin-containing state is present at steady state and could allow cells to permanently change secretion rates by adjusting the relative peflin expression level. On the other hand, the Ca^2+^-activated inhibitory state could “choke” secretion quickly during transient excitotoxic circumstances or viral infection. Though the consequences of secretion reduction could be protective to otherwise healthy cells, we speculated that it may be maladaptive to highly stressed cells. Along these lines we depleted peflin in a cellular model of ageing wherein primary porcine aorta endothelial cells become senescent, display mitochondrial Ca^2+^ excitotoxicity, elevated ROS production, chronic ER stress signaling in the absence of chemical inducers, and eventual apoptosis (31). We found that peflin depletion drastically reduced pro-apoptotic UPR signaling (Figure 5E), consistent with its suppressive role in secretion being relevant to damage-inducing ER stress.

## EXPERIMENTAL PROCEDURES

### Antibody Production and Puri/ication

Rat peflin and mouse ALG-2 were ligated into pGEX expression plasmids and expressed in E.coli as GST fusion proteins. Cultures were grown at 37°C to an A_600_ of 0.4-0.6, prior to an induction with 1mM isopropyl-1-thio-β-D-galactopyranoside (IPTG) at 37°C for GST-ALG-2, and 15°C for GST-peflin, for 3 h. Harvested cells were subjected to a single round of French Press and centrifuged at 20,000 x g for 20 min. Pellets were collected, dissolved in sample buffer and loaded onto SDS-PAGE gels. Gels were stained with 0.1% Coomassie in H_2_0, and the resolved bands were excised from the gel and subjected to a 3 h electroelution in 25mM Tris, 191mM glycine and 0.1% SDS, on ice. The eluted protein solution was concentrated and injected subcutaneously, using Freund’s adjuvant, into a rabbit, for peflin, or a chicken for ALG-2. Three subsequent antigen boost injections were done over an 80-day period. At this stage, the peflin antibody was fully useful as a crude serum. For the ALG-2 antibody, sera were supplemented with an equal volume of 10mM Tris, pH 7.5, filtered with a syringe filter and passed through a 1ml CNBr-Sepharose column conjugated with GST as non-specific control. The flow through was then loaded onto another CNBr-Sepharose column conjugated with mouse GST-ALG-2. Columns were washed with 3 x 5ml of 10mM Tris, pH 7.5, then washed with the same buffer containing 0.5M NaCl, once again with 10mM Tris, pH 7.5 and finally eluted with 0.1M glycine, pH 2.5. Fractions were neutralized with 2M Tris, pH 8.0, and quantitated at A280. Peak fractions were pooled and dialyzed into PBS.

### Other antibodies and expression constructs

Monoclonal anti-VSV-G was purchased from Sigma (St. Louis, MO: product V5507, clone P5D4), while monoclonal anti-VSV-G clone I14 antibodies (38) were produced in-house from the hybridoma cell line. Mouse monoclonal anti-CHOP antibody was purchased from ThermoFisher Scientific, Waltham, MA (product MA1-250). Rabbit polyclonal anti-collagen I antibody was purchased from Abcam, Cambridge, UK (product ab34710). Mouse monoclonal anti-mannosidase II antibody was purchased from Covance Research Products, Denver, PA (product MMS-110R-200). Green secondary antibodies conjugated to Alexa Fluor 488 were from Invitrogen (Carlsbad, CA: product A11001); Cy3-, or cy5-conjugated secondary antibodies were purchased from Jackson ImmunoResearch Laboratories (West Grove, PA).

Cargo constructs: for routine ER-to-Golgi transport studies, we used the synchronizeable cargo VSV-G_ts045_-GFP in pCMV, or more commonly the untagged version of VSV-G_ts045_ in pCMV in conjunction with the P5D4 monoclonal antibody for immunofluorescence detection (the untagged construct has a significantly higher transfection efficiency). Human GFP-Collagen I was from David Stephens via Addgene, Cambridge, MA (construct: pEGFP-N2-COL1A1). The transport cargos retained in the ER until triggered to export with a ligand are based upon the RPD Regulated Secretion/Aggregation Kit from ARIAD Pharmaceuticals. The luminal cargo we here call “GFP-F_M_4-GH” is identical to the construct “pC4S1-eGFP-F_M_4-FCS-hGH” we described before (24). GFP-F_M_4-VSV-G_tm_ was constructed from GFP-F_M_4-GH by removing the furin cleavage site and human growth hormone by cleavage with SpeI/BamHI and replacing it with a fragment containing the VSV-G transmembrane domain: 5′-actagtTCATCGTCGAAGAGCTCTATTGCCTCTTTTTTCTTTATCATAGGGTTAATCATTG GACTATTCTTGGTTCTCCGAGTTGGTATTTATCTTTGCATTAAATTAAAGCACACCAA GAAAAGACAGATTTATACAGACATAGAGATGAACCGACTTGGAAAGTAAGCGCCCG Cggatcc-3′. To make GFP-F_M_4-GPT was the same procedure except that the SpeT/BamHT-cleaved construct was ligated with a fragment containing the CD55 GPT anchor sequence: 5-actagtACAACCCCAAATAAAGGAAGTGGAACCACTTCAGGTACTACCCGTCTTCTATC TGGGCACACGTGTTTCACGTTGACAGGTTTGCTTGGGACGCTAGTAACCATGGGCTT GCTGACTTAGggatcc-3′. This construct is targeted to the plasma membrane where it is sensitive to extracellular PT-PLC treatment (DEG and AAP, unpublished observations). For VSV-G_ts045_ employed in Figure 7 as an ERES marker, we utilized the untagged VSV-G_ts045_, then immuno-labeled it using monoclonal antibody T14, which only detects mature trimers.

Rat peflin was amplified from a cDNA clone by PCR primers encoding EcoR1/XhoT. This product was ligated into mammalian expression vector pCDNA 3.1(+). Mouse ALG-2 was amplified from a cDNA clone (MGC: 49479) and ligated into mammalian expression vector pME18S using PCR primers encoding XhoT and XbaT. GFP-Sec13 was as described (39). Histamine receptor construct pH1R-P2A-mCherry-N1 was purchased from Addgene (product: 84330). For a plasma membrane marker for use in total cell fluorescence calculations, pCAG-mGFP was purchased from Addgene (product: 14757).

### siRNA knockdowns and trans/ections

For plasmid transfections, NRK or PC12 cells were transfected using Polyjet (SignaGen Laboratories; Frederick, MD), following the manufacturer’s instructions, about 24 h prior to transport assays. For transport experiments involving both plasmid and siRNA transfections, NRK or PC12 cells were transfected with siRNAs using RNAiMax (Tnvitrogen; Carlsbad, CA) with OpiMEM medium as suggested by the manufacturer, approximately 48 h prior to transport. Then, approximately 24 h prior to transport, they were transfected with plasmids using Polyjet as just described. Cells were equilibrated at 41 oC for 6-12 h prior to transport. Control siRNA had the following sense strand sequence: 5′-AGGUAGUGUAAUCGCCUUGdTdT-3′. Peflin siRNA 0975 had the following sense strand sequence: 5′-GCCUCAUGAUGAUAAACAU-3′, and consistently achieved ~95% knockdown in NRK cells as assayed by immunoblotting. Tn a previous manuscript, we established that three distinct, non-overlapping peflin siRNA sequences including 0975 - produced the phenotype of elevated ER-to-Golgi transport ((16), Figure S1). ALG-2 siRNA 8567 had the following sense strand sequence: 5′-GGAGCGGAGUGAUUUCAGA-3′ and consistently achieved ~95% knockdown. In a previous manuscript, we established that three distinct, non-overlapping ALG-2 siRNAs - including 8567 produced consistent, mild effects on ER-to-Golgi transport (8). Immunoblotting of cell lysates enabled validation of knockdown efficiencies for each siRNA experiment that was functionally analyzed. Collagen siRNA had the following sense strand sequence: 5′-GAACUCAACCUAAAUUAAAdTdT-3′. All siRNAs were custom synthesized lacking chemical modifications by Gene Link (Elmsford, NY).

### Cell culture and agonist/drug treatments

NRK and Rat2 cells were grown in DMEM with 4.5 g/L glucose, 10% FBS (Gibco, qualified grade), and 1% penicillin-streptomycin. For microscopy studies, they were grown directly on glass coverslips coated with poly-L-lysine. PC12 cells were grown in DMEM with 4.5 g/L glucose, 5% donor horse serum (Hyclone) and 5% iron-supplemented bovine calf serum (Hyclone). PC12 cells were maintained routinely on collagen-I-coated plasticware, while for microscopy they were plated directly on glass coverslips that had been first coated with poly-L-lysine and then coated with collagen I. Porcine aorta endothelial cells were isolated and cultured to P5 as described before (31). For agonist/drug treatments NRK or PC12 cells were grown to ~100% confluency in 6-well plates. Histamine (Sigma H7125) was dispensed into glass vials under nitrogen and stored at −20 °C. Solutions of histamine were prepared fresh each day, while ATP (500 mM in water) and BHQ (100 mM in DMSO) stock solutions were stored frozen and freshly diluted each day.

### PAEC’s and RT-PCR

Porcine aorta endothelial cells (PAECs) in an aged state (passage 5), confirmed by positive beta-galactosidase staining and decreased proliferation rate (31) were transfected with peflin siRNA using Transfast (Promega Corp., Madison WI, USA) according to manufacturer’s instructions. Total RNA was isolated using the PEQLAB total RNA isolation kit (Peqlab; Erlangen, Germany) and reverse transcription was performed in a thermal cycler (Peqlab) using a cDNA synthesis kit (Applied Biosystems; Foster City, CA). mRNA levels were examined by qRT-PCR. A QuantiFast SYBR Green RT-PCR kit (Qiagen; Hilden, Germany) was used to perform real time PCR on a LightCycler 480 (Roche Diagnostics; Vienna, Austria), and data were analyzed by the REST Software (Qiagen). Relative expression of specific genes was normalized to porcine GAPDH as a housekeeping gene. Primers for qRT-PCR were obtained from Invitrogen (Vienna, Austria).

### Calcium Imaging

Prior to imaging, NRK or PC12 cells were loaded by incubation in growth medium containing 20 mM Hepes, 3 μM FURA-2AM and 1.5 mM probenecid for 30 minutes at 37 °C, then washed and incubated for 10 min in the same medium lacking FURA-2AM. Coverslips were then placed in a microscope chamber and maintained at 37 °C using a submerged heating loop. For ATP experiments, cells were maintained in a non-perfusing volume of 1.5 ml growth medium, 20 mM HEPES and probenecid. For BHQ experiments, the medium was continuously perfused with pre-warmed medium at 2 ml/min. Image acquisition was completed by a Nikon TE300 inverted microscope equipped with a Nikon Plan Fluor 20x/0.75 objective, motorized high speed Sutter Lambda filter wheel for emissions, CoolLED pe340 excitation system, and PCO Panda sCMOS camera, all automated with Micro-Manager software. After selecting a suitable field of cells, imaging was carried out for at most 30 min of 10-second imaging cycles capturing separate 340 nm- and 380 nm-excited images collected at 510 nm. Analysis was completed in ImageJ using a custom plugin that determined the ratio of the background-subtracted emissions from 340 and 380 excitation for each individual cell over time.

### Immunofluorescence microscopy

Coverslips were fixed with 4% paraformaldehyde containing 0.1 M sodium phosphate (pH 7) for 30 min at room temperature and quenched three times for 10 min with PBS containing 0.1M glycine. Fixed cells were treated for 15 min at room temperature with permeabilization solution containing 0.4% saponin, 1% BSA, and 2% normal goat serum dissolved in PBS. The cells were then incubated with primary antibodies diluted in permeabilization solution for 1 h at room temperature. Next, coverslips were washed 3x with permeabilization solution and incubated 30 min at room temperature with different combinations of Alexa Fluor™ 488-, Cy3-, and/or Cy5- conjugated anti-mouse, anti-rabbit, or anti-chicken secondary antibodies. After the secondary antibody incubation, coverslips were again washed 3x using permeabilization solution and mounted on glass slides using Slow Fade Gold antifade reagent (Invitrogen: S36936) and the edges sealed with nail polish. Slides were analyzed using a 40x/1.3 Plan Fluor or 60x/1.4 Plan Apo objective on a Nikon E800 microscope with an LED illumination unit (CoolLED pE 300^white^), sCMOS PCO.edge 4.2 camera, Prior excitation and emission filter wheels and Z-drive, automated using Micro-Manager software. For transport assays (see below) typical images collected for each field of cells were VSV-G_ts045_ (green channel) and Golgi marker mannosidase II (cy5 channel). For colocalization assays (see below) typical images collected for each field of cells were Sec13-EGFP (GFP channel), ALG-2 (cy3 channel), and peflin (cy5 channel).

### ER-to-Golgi transport assay

NRK, PC12, or Rat2 cells were plated on glass coverslips and transfected as described above. For VSV-G_ts045_ and collagen I cargo, cells were shifted to 41 °C for 6-15 h prior to transport, to accumulate the cargo in the ER. For the transport assay, the cells were either fixed by dropping coverslips directly into 6-well chambers containing fixative or pre-equilibrated 32 °C medium for 10 min, then fixative. For assays involving collagen I cargo the 32 °C medium was supplemented with 50 μg/ml ascorbate. Alternatively, for F_M_4-containing cargo constructs, transfected cells were kept always at 37 °C, and the coverslips were fixed either by dropping coverslips directly into fixative or into to 6-well chambers containing 37 °C media with 500 nM AP21998, also known as D/D solubilizer (TakaraBio, Shiga Japan: 635054), for 10 min prior to transfer to fixative.

Morphological quantitation of ER-to-Golgi transport was accomplished by first collecting images in a consistent manner with regard to cell morphology, protein expression levels and exposure. A single widefield image plane was collected for each color channel for each field of cells randomly encountered; image deconvolution was not performed. Prior to image analysis using a custom ImageJ script (available upon request), files for all experimental conditions were automatically and randomly renamed with a 36-character designation and re-sorted by that identifier, eliminating any way for the user to know their identities. Each image is opened in turn and presented to the user, who at this point only views the cargo image plane. For each image the user first defines an extracellular region for use as a background, after which the user defines the minimal rectangular region of interest (ROI) encompassing the first cell to be analyzed. This ROI is then isolated in a separate window and the Golgi maximum is extracted, which represents the mean intensity of the pixels in the 99.990 percentile and above but excluding the highest pixel. The user checks that these brightest pixels are in fact within the Golgi as defined on the mannosidase II image planes. The user then sequentially places 3 small circular ROIs within vesicular/reticular regions adjacent to the nucleus but clearly distinct from the Golgi. The ER mean is extracted as the mean of the three mean pixel intensities of these ROIs. Transport index is then extracted for the cell as (Golgi maximum-background) / (ER mean-background). The cell is then numbered on the image to avoid re-counting, and all extracted parameters written to an appendable output file along with the cell number, and image title so that the data is fully traceable. The user then defines another cell from the image or opens another image. Using this method, the user quantitates 60-100 cells per hour.

Once transport indices have been obtained for all conditions in an experiment, each value is subtracted by the mean transport index value for cells that were fixed directly from 40 °C without a transport incubation at 32 °C (typically a value between 1.0 and 1.5) to generate the net transport index. Net transport indices are then normalized to the mean control value for the particular experiment. Each result reported here was obtained in at least three separate experiments on different days.

### Labeling intensity and colocalization assays

Cells were transfected with ERES markers GFP-sec13, or VSV-G_ts045_. Following immunolabeling as described above, cells such as those shown in Figures 2 and 7 (or quantitated in Figure 10E) were captured as z-stacks in 11 200-nm increments for each channel. These image stacks were deconvolved as a single batch using Huygens Essential Widefield software (Scientific Volume Imaging, Hilversum, The Netherlands) according to the manufacturer’s instructions. Final images for display and quantitation represent maximum intensity projections of deconvolved stacks. As before, these deconvolved stacks were each assigned a random 36-character designation and re-sorted by the random name. The intensity of labeled proteins was assessed by a custom ImageJ script (available upon request) performed on individual cells for which the user manually defines a minimal enclosing ROI. Background labeling was first removed by defining a dark extracellular area of each image channel as background, and subtracting that value from every pixel. The specific pixels assessed for intensity were predetermined using a binary object mask, which was generated by auto-thresholding the desired area of the cell using the Renyi Entropy or Intermodes algorithm, depending upon which most accurately captured the spots of interest (but kept constant for a given marker/protein). Spots on the mask were assessed as ROIs that were used to measure either mean intensity or integrated density (product of area and mean intensity) in relevant channels of the unmodified images. Integrated density measurements are referred to as “total intensity” or “total spot intensity” in the figure legends where relevant, whereas mean intensity measurements are labeled as such. The choice to present total vs. mean intensity was driven by which parameter produced lower variance and thus lower p values when used in T-tests (in no cases did they produce opposing results). In a few cases (Figure 7C and D), the binary mask used was actually the Boolean intersection (see below) of ALG-2 and another ERES marker (sec31A or peflin); this was because the ALG-2 antibody produced suspected background spots that did not co-localize with ERES markers and were not removed by ALG-2-specific siRNAs. In our nomenclature, “ERES intensity” and “spot intensity” are distinguished by whether the mask for interrogation is generated from a distinct ERES marker from that being measured or from the measured marker itself, respectively. Extracted parameters were written to an appendable output file along with the cell number and image title so that the data was traceable. Following re-sorting by experimental conditions, integrated densities and other parameters for each cell were then normalized to the mean control value for the particular experiment. Each result reported here represents combined data from at least three separate experiments that displayed similar trends.

Alternatively, some binary masks were used to assess particle areas or areas of overlap - also referred to as “co-localization”. Areas of overlap between two channels were calculated from two auto-thresholded images using the Boolean ‘AND’ operator in ImageJ’s image calculator. This operation generated a mask that contained only spots present in both channels, permitting a calculation of the total overlap area.

### Total Cell Fluorescence Assay

In Figures 5B and C, total cell fluorescence of collagen was determined by first transfecting Rat2 cells with the plasma membrane marker pCAG-mGFP (GFP with an N-terminal palmitoylation signal). Endogenous collagen I was labelled in the cy3 channel. Using pCAG-mGFP, a whole cell ROI was selected via the wand threshold tool in ImageJ. That ROI was then moved into the collagen channel wherein total mean gray values and ROI area were extracted. Separately, a mean background value was extracted by randomly selecting an area without a cell. Total cell fluorescence was calculated as area of selected cell x (mean intensity of cell - mean intensity of background).

## ACKNOWLEDGMENTS

This work was supported by NIH grant 1R15GM106323-02 to JCH, and the University of Montana Center for Biomolecular Structure and Dynamics NIH grant P20GM103546.

## Notes

### Competing Interest Statement

The authors have declared no competing interest.

### Summary of Updates

This revision adds new data that significantly strengthens the conclusions and provides a graphical model of PEF protein mechanisms.

