## Supplemental Figure 1 for "ALG-2 and Peflin Stimulate or Inffibit Copii Targeting and Secretion in Response to Calcium Signaling"

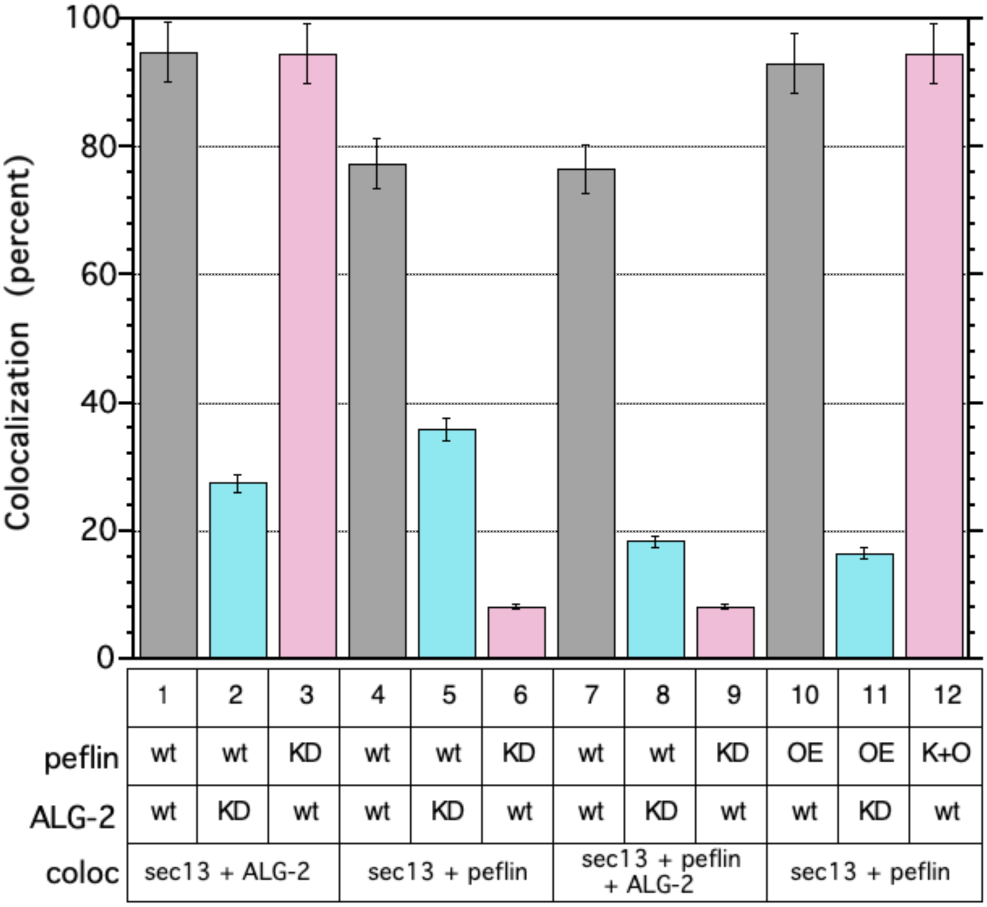


**Supplemental Figure 1. Manual co-localization analysis for the experiment shown in Figure 2.**

Methods: Deconvolved images from ~25 transfected cells containing in focus GFP-Sec13 labeling at ER-exit-sites were randomly selected, and the background labeling was removed by defining a dark extracellular area of the image as zero. A GFP-sec13 object binary image mask was generated in Fiji by auto-thresholding using the Renyi Entropy method, which consistently captured the brightest GFP-sec13 spots. Each of these objects were then tested by eye for co-localization with spots in the ALG-2 and/or peflin channel with overlap scored as positive for co-localization. The percentage of bright ALG-2 and/or peflin objects that overlapped with ERES spots was directly calculated from this analysis for each cell, and the values above represent means of ~25 cells per condition. Mean ± SEM is shown for each condition.
